# A Universal Stress Protein upregulated by hypoxia may contribute to chronic lung colonisation and intramacrophage survival in cystic fibrosis

**DOI:** 10.1101/2020.10.03.324806

**Authors:** Andrew O’Connor, Rita Berisio, Mary Lucey, Kirsten Schaffer, Siobhán McClean

**Author notes:** Corresponding author: Dr Siobhán McClean.

## Abstract

Universal stress proteins (USPs) are ubiquitously expressed in bacteria, plants and eukaryotes and play a lead role in adaptation to environmental conditions. In Gram negative bacteria they enable adaption of bacterial pathogens to the conditions encountered in the human niche, including hypoxia, oxidative stress, osmotic stress, nutrient deficiency or acid stress, thereby facilitating colonisation. We previously reported that all six USP proteins encoded within a low-oxygen responsive locus in *Burkholderia cenocepacia* showed increased abundance during chronic colonisation of the CF lung. However, the role of USPs in chronic infection is not known. Using mutants derived from *B. cenocepacia* strain, K56-2, we show that USP76 is required for growth and survival in many conditions associated with the CF lung including, hypoxia, acidic conditions, oxidative stress. Moreover, it is involved in attachment to host epithelial cells, but not virulence. It also has a role in survival in macrophages isolated from people with CF. In contrast, another USP encoded in the same locus, USP92 had no effect on host cell attachment or oxidative stress, but was responsible for a 3-fold increase in virulence. Overall this shows that these USPs, both upregulated during chronic infection, have distinct roles in *Burkholderia* pathogenesis and may support the survival of *B. cenocepacia* in the CF lung. Specifically, USP76 is involved in its survival within CF macrophages, a hallmark of *Burkholderia* infection.

## Introduction

Many opportunistic bacterial pathogens must adapt as they transition from their natural environment to the host in order to survive within the host and establish an infection. Changes in temperature, pH, osmolarity and oxygen availability are among the stresses that bacteria must overcome as they move from their environmental niche to that of mammals. Universal stress proteins (USPs) are expressed by bacteria and other microorganisms in response to a wide variety of environmental conditions. They are extensively expressed throughout nature from bacteria to archaea and eukaryotes but have not been identified in humans[1]. This breadth of evolutionary range illustrates the importance of USPs in the resilience of organisms to survive in stressful environments and while their role in bacterial cells is predominantly an adaptive response to the changing environment, their contributions to bacterial pathogenesis cover a range of different mechanisms [1]. In opportunistic pathogens such as *Mycobacterium tuberculosis, Staphylococcus aureus* and *Pseudomonas aeruginosa*, USPs are involved in survival within macrophages and low oxygen conditions [2–4], while an *Acinetobacter baumannii* USP has a protective role against low pH and contributes to pathogenesis [5].

USPs were first identified in *E.coli* K-12 following exposure of cells to a variety of stresses, heat shock, carbon and nitrogen starvation and ultraviolet radiation [6]. Expression of UspA in *E. coli* was independent of the stringent stress response transcriptional activators RelA/SpoT, RpoH, KatF, OmpR, AppY, Lrp, PhoB and H-NS [7]. *E. coli uspA* gene deletion mutants were defective in survival over prolonged periods of growth under stress conditions such as induced peroxide stress and osmotic shock [8]. *M. tuberculosis* USP Rv2623 contributes negatively to virulence by regulating its growth in the transition to latency [4]. There are six *usp* genes in the *E. coli* genome, while there are 10 *usp* genes encoded on the *M. tuberculosis* genome [1]. *Burkholderia cepacia* complex (Bcc) is a group of Gram negative bacteria that naturally occurs in soil and in the rhizosphere of crop plants and causes chronic opportunistic life-threatening infections in people with cystic fibrosis (CF) and immunocompromised patients. It can also colonise pharmaceutical plants and contaminate pharmaceutical products and disinfectants [9]. It is highly antimicrobial resistant and once a chronic infection has been established, eradication is rare. Its capacity to colonise diverse and harsh niches are exemplary and consequently elucidation of its mechanisms of adaptation is essential. Sass *et al.* (2013) identified a 50 gene locus which was dramatically upregulated under low oxygen conditions and designated the low oxygen activated (lxa) locus [10]. We subsequently showed that 19 proteins encoded on the lxa locus showed increased abundance in late infection isolates from chronically colonised CF patients [11]. These late chronic infection isolates also showed increased attachment to CF lung epithelial cells [12]. Importantly, all six USPs encoded on the lxa locus showed increased abundance with time of colonisation [11]. Among these lxa-encoded *usp* genes, BCAM0276, encodes a UspA family stress protein which consistently showed increased abundance in the later isolates from two chronically colonised patients and was associated with increased gene expression [11]. Previously this USP (USP76) was reported to be upregulated almost 60-fold in *B. cenocepacia* strain J2315 response to low oxygen [10] and up to 40-fold in a comparative transcriptomic study of *B. contaminans* isolates from CF patients [13]. There are 11 USPs in total encoded on the *B. cenocepacia* genome [14], ten of which are on chromosome 2. The role of USP76 or other USPs in Bcc are unknown, but this increased expression during chronic infection and under low oxygen conditions suggests that this gene may be involved in chronic persistence of *Burkholderia cepacia* complex during infection of the hypoxic CF lung. While USPs have been shown to protect bacteria from a range of environmental pressures and stresses including extreme temperature changes, antibiotic challenges, nutrient deprivation and oxidative stress, to date there are no published data on the roles of USPs in chronic infection.

Limited oxygen is a hallmark of the CF lung due to rapid oxygen consumption by microorganisms, invading neutrophils, impaired ventilation and contributions from mucous plugs which together create an oxygen gradient within the lung [17]. These selective pressures in CF airways drive adaptation in colonising bacteria, enabling them to overcome the various microenvironments within the CF lung during infection. Furthermore, pathogens such as Bcc can survive phagocytosis and even replicate within macrophage contributing to chronic infection [18, 19]. We derived two *Δusp* deletion mutants and demonstrate that USPs such as USP76 are at the forefront of this adaptation and may play a role in *B. cenocepacia* pathogenesis. In addition, we compare it to another USP encoded on the lxa locus, which has a comparable predicted size (table 1) and was also increased in abundance over time of chronic infection and show the two USPs are distinct.

**Table 1:**
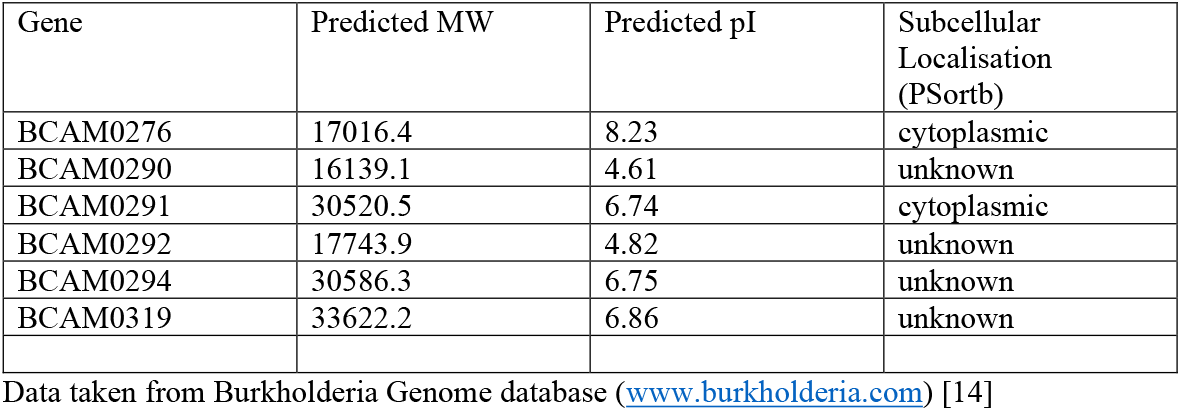
USPs encoded on the lxa locus in *B. cenocepacia* J2315.

| Gene | Predicted MW | Predicted pI | Subcellular Localisation (PSortb) |
| --- | --- | --- | --- |
| BCAM0276 | 17016.4 | 8.23 | cytoplasmic |
| BCAM0290 | 16139.1 | 4.61 | unknown |
| BCAM0291 | 30520.5 | 6.74 | cytoplasmic |
| BCAM0292 | 17743.9 | 4.82 | unknown |
| BCAM0294 | 30586.3 | 6.75 | unknown |
| BCAM0319 | 33622.2 | 6.86 | unknown |

Data taken from Burkholderia Genome database (www.burkholderia.com) [14]

## Results

USP76 and USP92 both belong to the UspA protein family (PF00582) and contain single UspA domain from residue 1 to 145. Both are predicted to exist as a homodimer formed largely by the C-terminus of each monomer based on Swiss-Model Expasy modelling (Figure. 1A), which is common among other UspA molecules [20]. The BCAM0276 gene is the one of six USPs encoded on the lxa locus. USP76 showed 6-fold increased abundance from early to late infection in both sets of sequential isolates and was selected for detailed investigation.

**Figure 1:**
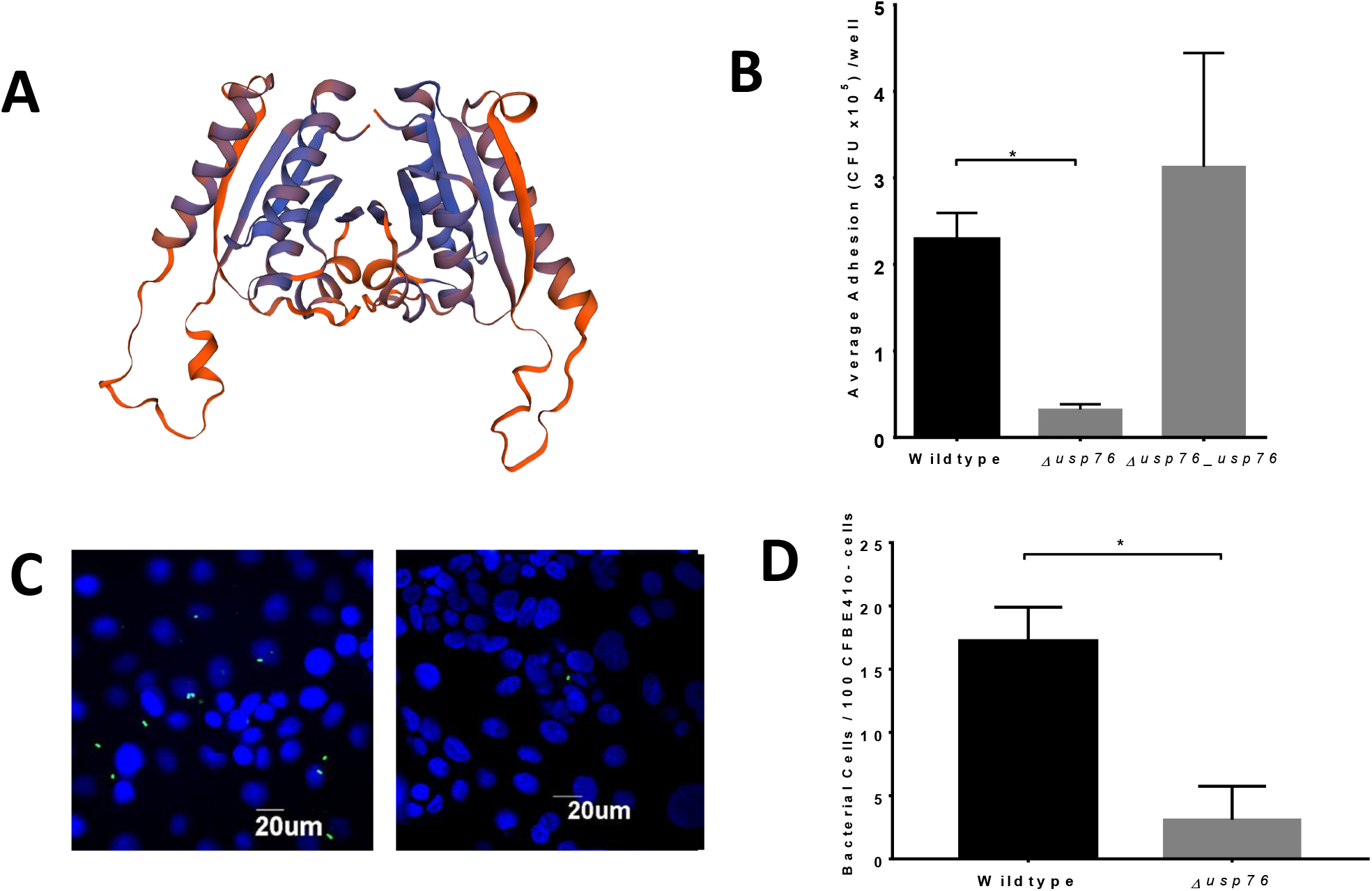
Comparison of the bacterial attachment to CFBE41o^-^ cells measured by microbiological plating at an MOI of 50:1. **A)** Predicted structure of the USP76 homodimer using Swiss-Model Expasy. b) Comparison of the attachment of WT K56-2 and the *Δusp76* mutant to CFBE41o^-^ lung epithelial cells by (b) microbiological plating and (c & d) confocal microscopy. B) Data represent the mean CFU / ml for each strain in three independent experiments. Error bars represent the standard error of the mean. *Signifies a statistically significant difference in attachment of the *Δusp76* mutant as determined by one-way ANOVA, p = 0.0117. C) Confocal microscopy images of the attachment of WT K56-2 and the *Δusp76* mutant to CFBE41o^-^ cells at a MOI of 50:1. *B. cenocepacia* cells were labelled with a primary anti-Bcc antibody and detected with a secondary FITC-conjugated antibody (green). CFBE nuclei were counterstained with DAPI (blue). D) Data represents the mean number of bacteria / 100 cells of CFBE41o^-^ per strain for 10 randomly selected fields of view in three independent experiments. Error bars represent the standard error of the mean. * Signifies statistical significance, using a t test to compare data from three independent experiments, *p < 0.0001.

### The *Δusp76* mutant has a reduced attachment to host cells

We previously showed that the attachment of sequential *B. cenocepacia* isolates to CFBE41o^-^ cells increased over the time of chronic infection [12]. Given the observed increase in BCAM0276 gene expression and corresponding increase in USP76 protein abundance [11], the effect of the deletion of BCAM0276 in *B. cenocepacia* strain K56-2 on attachment to cystic fibrosis epithelial cells, CFBE41o^-^ was examined. The resulting *Δusp76* mutant showed 90% reduced attachment to CFBE41o^-^ cells (p < 0.05)(Figure 1b), which was restored to wildtype levels in the *Δusp76_usp76* complement strain. The reduction in attachment of the *Δusp76* mutant was also confirmed by confocal microscopy (a 6-fold reduction in attached *Δusp76* mutant cells compared to WT (p < 0.05, Figure 1 c and d)

### The *Δusp76* mutant elicits reduced cytokine response but does not impact on acute virulence

Given that deletion of the BCAM0276 gene resulted in reduced attachment to CFBE41o^-^ cells, the secretion of IL-8 and IL-6 cytokines by CFBE41o^-^ cells was examined to investigate whether there was a concomitant effect on cytokine secretion. IL-8 secretion was impaired in response to the *Δusp76* mutant strain (Figure 2a; p = 0.0135) while no statistically significant effect on IL-6 relative to wild-type was observed. Despite the reduced IL-8 response of the *Δusp76* mutant, both the WT and the mutant strains were highly virulent, with 0% survival at all dilutions of inoculum at 48 hours and consequently the LD_50_ was determined at 24 hours (Figure 2c). The *Δusp76* mutant strain showed comparable virulence in the *G. mellonella* acute virulence infection model.

**Figure 2:**
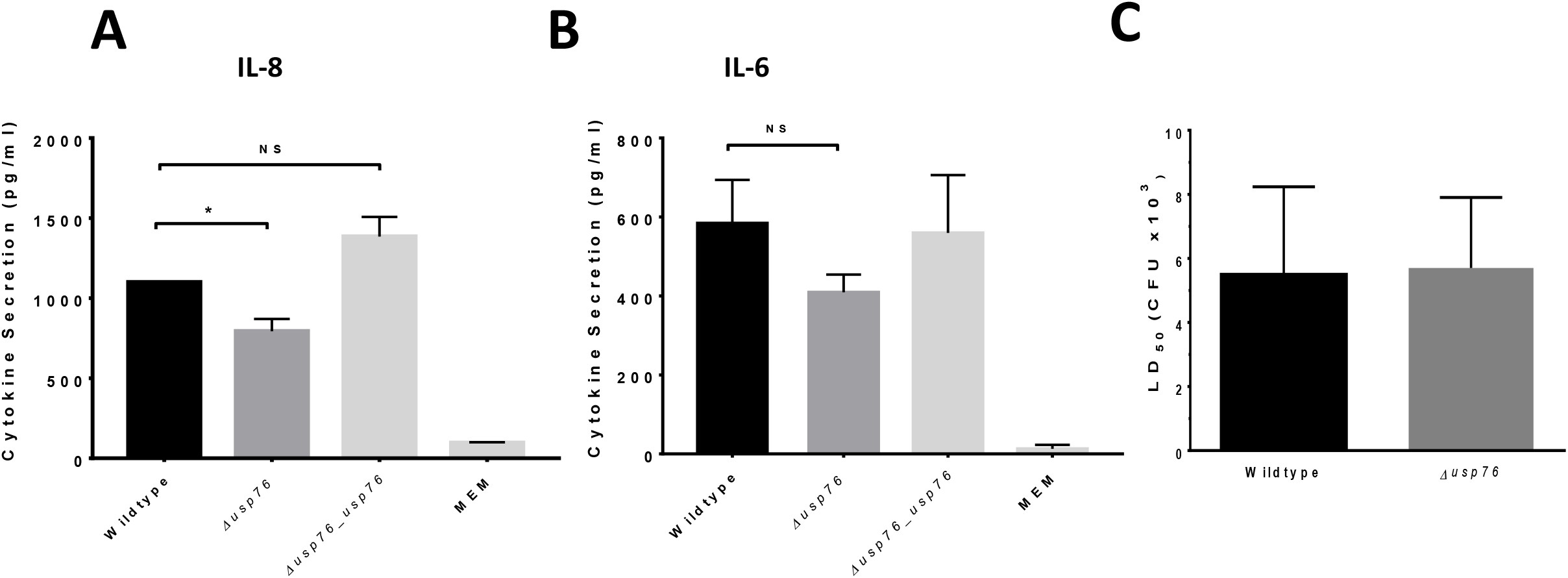
Effect of deletion of BCAM0276 on cytokine response and virulence. A & B) IL-8 and IL-6 cytokine secretion from CFBE41o^-^ cells following infection with the WT, the *Δusp76* mutant, the *Δusp76_μsp76* complement strains or negative control (MEM only). Bars represent the mean level of detected cytokine in triplicate measured in two independent experiments. Error bars represent the standard deviation. Statistical analysis was performed using one-way ANOVA, *p = 0.0135**. C)** LD_50_ values for the WT and the *Δusp76* mutant strains in the *G. mellonella* acute infection model at 24 hours. Data represent the mean LD_50_ from three independent experiments and error bars represent the standard error of the mean.

### USP76 is required for survival in low oxygen and under an oxidative environment

Previous studies on bacterial USPs highlight roles in survival during challenging environmental conditions [1], given the increase in BCAM0276 gene expression and in abundance of USP76, the response of *Δusp76* under various environmental pressures experienced during chronic infection was examined. It has been shown that the lxa locus, including the BCAM0276 gene, was dramatically upregulated in response to low oxygen [10] therefore it was important to understand if USP76 was involved in survival in response to low oxygen. The K56-2 strain was unable to grow in oxygen environments lower than 6% oxygen and survival was considerably impaired at 6% oxygen (Figure 3). When exposed to 6% oxygen in a controlled hypoxia chamber, no *Δusp76* mutant cells survived to day 8, indicating that this gene is critical to survival under low oxygen. Complementation of the gene in the *Δusp76_usp76* strain restored survival to near WT levels.

**Figure 3:**
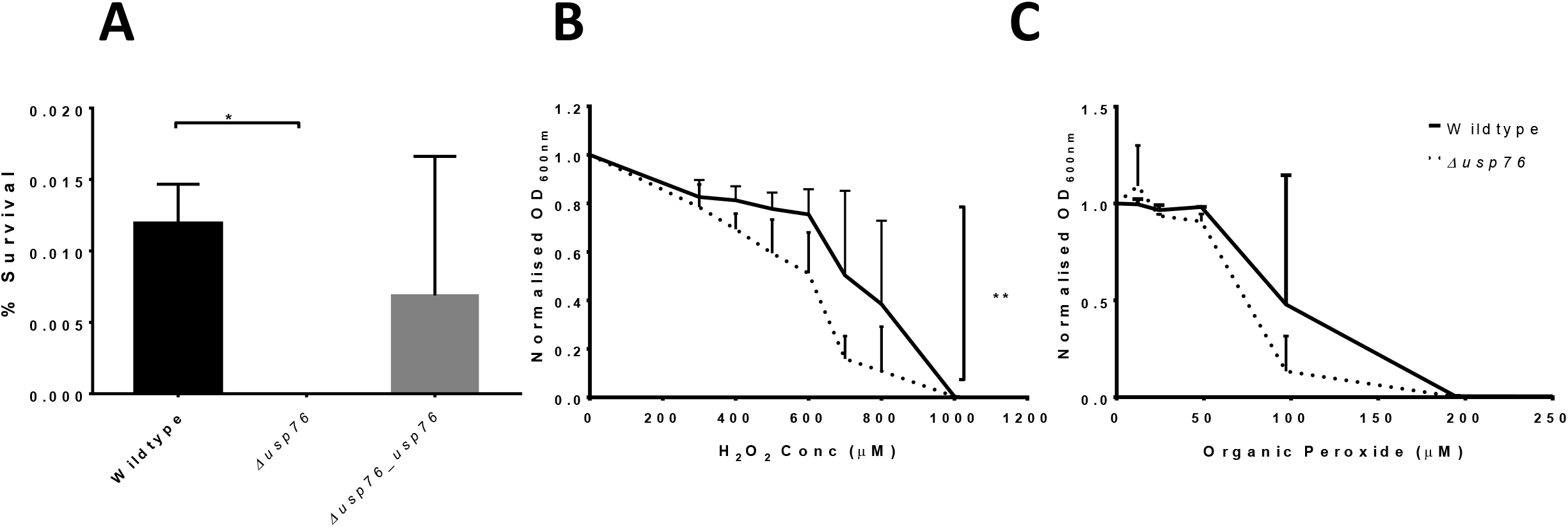
Effect of USP76 on survival or growth to environmental conditions. **A)** Mean percentage survival of the WT, the *Δusp76* mutant and the *Δusp76_usp76* complement strains after 8 days incubation in 6% oxygen in a controlled hypoxia chamber determined in three independent experiments. Error bars represent the standard error of the mean; *p = 0.0258. **B & C)** Mean OD_600_ values of the WT and *Δusp76* mutant strains following incubation with a range of concentrations of **B)** H_2_O_2_ or **C)** Tert-butyl hydroperoxide (0 - 1000 μM) for 20 hours. Data displayed was normalised relative to treatment-free control and represent duplicate values of three independent experiments, ** p=0.0016.

The CF lung is also characterised by constant inflammation and production of reactive oxygen species (ROS) due to persistent bacterial infections. The impact of the oxidative environment on the survival of the *Δusp76* mutant was examined using inorganic peroxide (H_2_O_2_) and organic peroxide (tert-butyl hydroperoxide). Growth of the *Δusp76* mutant was impaired when cultured in the presence of hydrogen peroxide (p = 0.0016, Figure 3b) although no significant difference in growth was observed for tert-butyl hydroperoxide over a period of 20 hours (Figure 3c, p = 0.2731).

### USP76 is required for growth and survival at acidic pH and survival in nutrient limited conditions

A number of bacterial USPs allow survival under acidic conditions. Growth of the *Δusp76* mutant in LB at low pH (pH 4.5) was reduced (p < 0.003) over 20 hours relative to WT when normalised for growth in LB at physiological pH (~pH 7.4, Figure 4a). Growth of the *Δusp76_usp76* complemented strain was comparable to WT levels. When the strains were plated to determine the viable CFU remaining after exposure to acidic pH at a range of time points, it was apparent that viability of the K56-2 strain following growth in acidic conditions was also dependent on USP76 expression (Figure 4b), particularly after 2 hours exposure (p = 0.0092). The *Δusp76_usp76* complement strain initially showed impairment of viability, but this recovered over time to reach WT levels within 5 hours of exposure to acidic pH.

**Figure 4:**
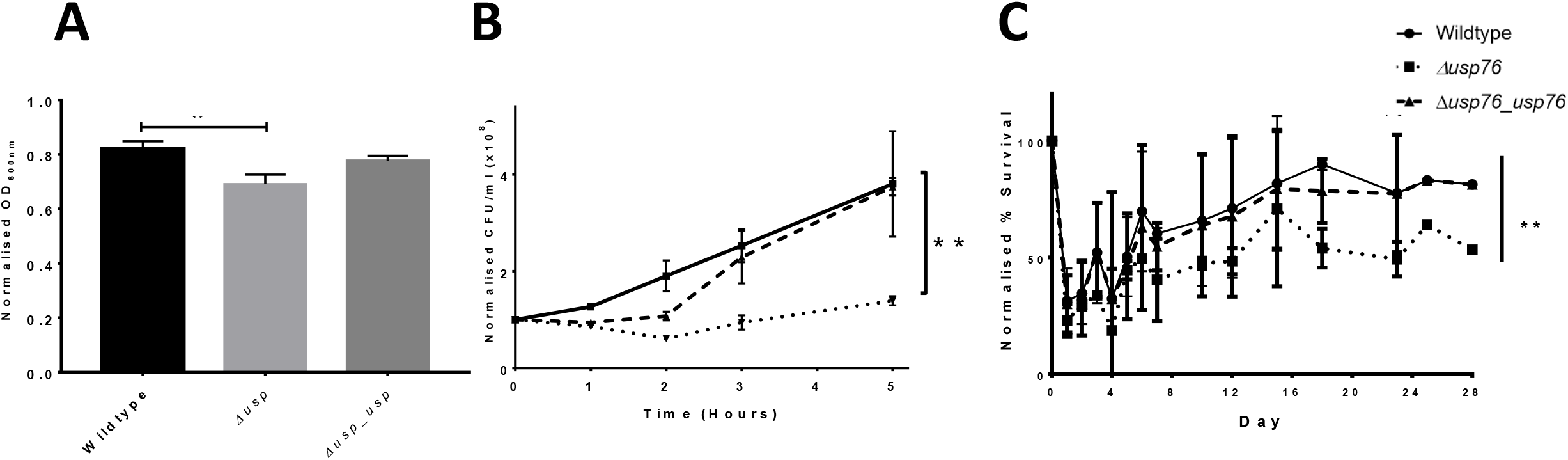
Growth and Survival of the *Δusp76* mutant under induced pH 4.5 stress or nutrient limiting conditions. **A)** Mean normalised endpoint OD_600nm_ values for each strain WT, the *Δusp76* mutant and *Δusp76_usp76* complement following incubation in pH 4.5 LB versus LB medium at pH 7. Data represent the mean normalised absorbance from 3 independent experiments, and error bars represent the standard error. Statistical analysis was performed by t test, **p = 0.0068. B) Normalised survival (CFU/ml) over time for each strain the *Δusp76* mutant and the Δ*usp76_usp76* complement strains compared to T_0_ following incubation in LB pH 4.5. Data represent the mean normalised CFU/ml from 3 independent experiments, error bars represent the standard error. Statistical analysis by two-way ANOVA, **p = 0.0019. C) Mean % survival of the WT, the *Δusp76* mutant and the *Δusp76_usp76* normalised to Day 0 of each respective strain following incubation in nutrient limiting medium (glucose free) over the course of 28 days. Data represent the mean % survival of 2 independent experiments. Statistical analysis performed by two-way ANOVA, p = 0.0038

The effect of deletion of the BCAM0276 gene on survival under long term nutrient limited conditions was also examined over 28 days. All three strains showed reduced survival over the first 3 days in glucose free medium, however both the WT and the *Δusp76_usp76* complemented strains recovered within 5 days, reaching almost 100% survival within 20 days. In contrast, the *Δusp76* mutant strain did not recover to the same extent, reaching 50 % survival by day 28 (p =0.0001, Figure 4c).

### Impact of USP76 on cellular permeability and antibiotic susceptibility

The *Δusp76* mutation had a moderate effect on cellular permeability as determined by the uptake of the cell permeable Hoechst dye over a two-hour period (p < 0.0001; Figure S1). The *Δusp76_usp76* complement showed much higher fluorescent intensity than the WT and *Δusp76* strains, possibly due to the deletion of the BCAL1674-1675 encoded efflux pump during the complementation process, which resulted in greater accumulation of Hoechst 33324 dye. In contrast to the alteration in cellular permeability, the BCAM0276 gene did not have any role in survival in response to antibiotic challenges encountered during treatment for CF infection. Growth following exposure to antibiotics with a range of mode of actions (Meropenem, Levofloxacin, Ceftazidime and Polymyxin B) was comparable for the WT and the *Δusp76* mutant strains (p = 0.909) (Figure S2, S3). The CF lung and the airway surface liquid also represent high salt environments with 1% w/v NaCl concentrations reported for the ASL of people with CF compared to 0.7% in healthy controls [21]. This creates osmotic stress on any pathogens colonising the CF lung. Growth of the *Δusp76* mutant was comparable to that of the WT strain over a range of salt concentrations (0 to 5% NaCl, p = 0.72) indicating that USP76 does not play a role in the salt tolerance of *B. cenocepacia* (Figure S4).

### Deletion of *usp76* has no effect on mucoidy, motility or biofilm formation

Alterations in mucoid phenotype have been associated with chronically infecting Bcc isolates [22], however, deletion of the *usp76* gene did not impact on EPS production when grown on YEM agar plates in either normoxic or hypoxic conditions (Figure S5, Table S3). Motility was comparable between the WT and the *Δusp76* mutant strain (Figure S6). No statistically significant differences were observed for motility phenotype when analysed by ANOVA, swimming p = 0.9720, swarming p = 0.2044 and twitching p = 0.4226. In addition, the ability for *B*. *cenocepacia* to form biofilms was not affected by the presence of the *usp76* gene (Figure S7) under normal oxygen conditions or hypoxic conditions.

### USP76 is required for survival of *B. cenocepacia* in CF macrophages

Bcc can survive and replicate within macrophage [18] allowing the organism to evade both host immune response and antibiotic treatments. Given that the pH of the intramacrophage phagosome is estimated to be between 6.2 and 4.5, and that we have shown that the *Δusp76* mutant is more susceptible to low pH and oxidative induced stress, we wanted to investigate whether USP76 might contribute to the survival of *B. cenocepacia* inside macrophages. Firstly we examined whether the intracellular uptake into the U397 macrophage-like cell line was altered by the presence or absence of USP76 and found that there was a 60% reduction in the number of the *Δusp76* mutant cells that are internalised by the U937 line compared to WT (Figure 5a, p = 0.002). Phagocytosis of the *Δusp76_usp76* complement strain was equivalent to that of WT. Survival of the *Δusp76* mutant within U937 macrophages cells was also significantly impaired at 24 h (Figure 5b, p = 0.0378). In contrast survival of the complemented strain was restored to wild type levels. Acidification is impaired in CF macrophages due to the dysfunctional CFTR, consequently, in order to examine whether this had relevance in the CF context, we sought ethics approval to investigate the survival of the strains in PBMC-derived macrophages from 12 people with CF. PBMC samples from five subjects could be successfully differentiated to macrophage cells in sufficient numbers to be suitable for use in uptake and intracellular assays. The uptake of the *Δusp76* mutant strain by PBMC-derived macrophage from two patients was significantly reduced (**p = 0.0018, ***p = 0.0004) when compared to the WT strain. More importantly, intracellular survival of the *Δusp76* mutant was assessed in CF patient derived macrophage cells, and showed significantly reduced survival in four out of five macrophage preparations (Figure 6b) (0.05 < p > 0.0001).

**Figure 5:**
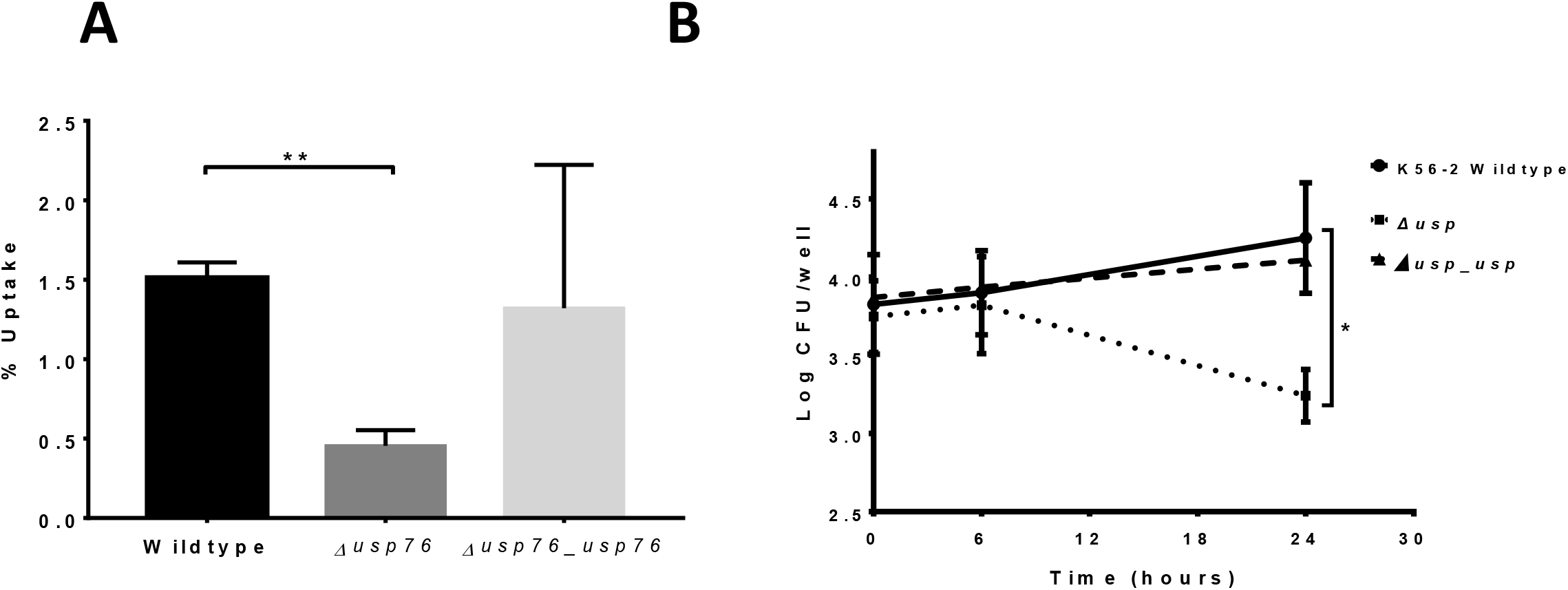
Uptake and survival of WT, the *Δusp76* mutant and the *Δusp76_usp76* complement strains in U937 macrophage cells. **A)** Uptake of the WT, the *Δusp76* mutant and the *Δusp76_usp76* complement strains by differentiated U937 macrophage cells. Data represent the intracellular uptake of the individual strains as a % of bacterial cells applied in three independent experiments. Error bars represent the standard error of the mean. **Statistically significant difference relative to the WT as determined by student t-test, p = 0.002); b) Survival of the *Δusp76* mutant and the *Δusp76_usp76* complement strains in U937 macrophage cells over time. Data represent the mean log_10_ of CFU/ml from three independent experiments; error bars represent the standard error of the mean. *Statistically significant difference relative to the WT determined by two-way ANOVA (p = 0.0378).

**Figure 6:**
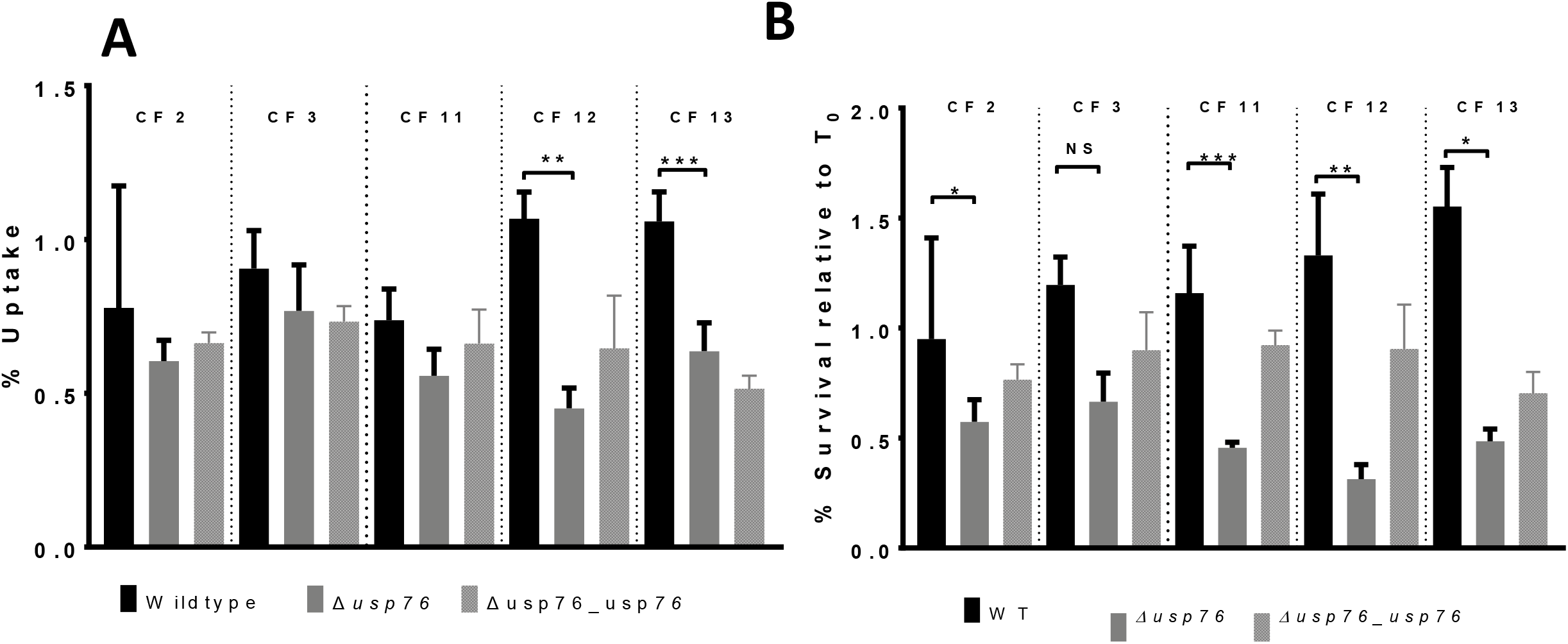
Uptake and survival of the WT, the *Δusp76* mutant and the *Δusp76_usp76* strains in PBMC-derived macrophage from people with CF. a) % uptake and b) 24 h survival of the WT (black), the *Δusp76* mutant (grey bars) or the *Δusp76_usp76* complement strain (grey patterned bars). The data represent the means of three independent experiments and error bars represent the standard error of each mean. Statistically significant difference relative to the WT as determined by one-way ANOVA (**p = 0.0018; ***p=0.0004). b) Survival of the wild type, *Δusp76* mutant and the *Δusp76_usp76* complement strains in CF monocyte-derived macrophage cells. Normalised data represent the mean log of CFU/ml from three independent experiments, relative to time zero. Error bars represent the standard error of the mean. Statistically significant difference between mutant and WT strains was determined by one-way ANOVA as follows: CF2: p = 0.0403; CF 11, p = 0.0016; CF 12 p = 0.0025; CF13, p<0.0001.

### The USPs encoded in the lxa Locus are not redundant

As previously mentioned there are six USPs encoded on the lxa locus, all of which showed increased abundance in late stage isolates from chronically colonised people with CF. Having characterised USP76, we wanted to examine whether other USPs expressed within the locus share the same functions. We chose a *uspA* gene downstream of BCAM0276 as it was upregulated by 201-fold in response to low oxygen and showed greatest increase in protein abundance (up to 16 fold) in late stage isolates [10, 11]. A targeted single deletion mutant was generated for the BCAM0292 gene; however, despite many attempts we were unable to complement the strain with the BCAM0292 gene. Using the targeted deletion mutant, *Δusp92*, the role of USP92 was characterised and compared with USP76. Consistent with *Δusp76*, the *Δusp92* mutant showed impaired growth under acidic conditions, relative to the WT strain K56-2 (Figure 7, p < 0.001). But in contrast to the *Δusp76* mutant, the *Δusp92* mutant showed a three-fold reduction in virulence as determined by LD_50_ in *G. mellonella* larvae relative to the WT strain (p = 0.0228, Figure 7). Moreover, the *Δusp92* mutant showed slower growth in the presence of high salt (Figure 7c p < 0.001) in contrast to *Δusp76*. USP92 may also confer slight but significant protection against sucrose at 2.5% (p = 0.04) and 5% w/v sucrose (p = 0.005)(Figure S8a). Furthermore, the *Δusp92* mutant also showed a slight but significantly reduced uptake of the cell-permeable Hoechst dye (p < 0.001). In contrast to the *Δusp76* mutant, no significant change in CBE41o^-^ cell attachment was observed in the *Δusp92* mutant compared with the WT strain (Figure S8b). Moreover, while USP76 seemed to be involved in survival in response to oxidative stress, the USP92 does not, with the *Δusp92* mutant showing comparable growth in the presence of hydrogen peroxide (Figure S8c). Consistent with the *Δusp76* mutant, there was no alteration in EPS production in the *Δusp92* mutant under normoxic conditions when compared to the WT strain (Figure S9a) nor were there significant difference in antibiotic susceptibility observed (Figure S9 b,c).

**Figure 7:**
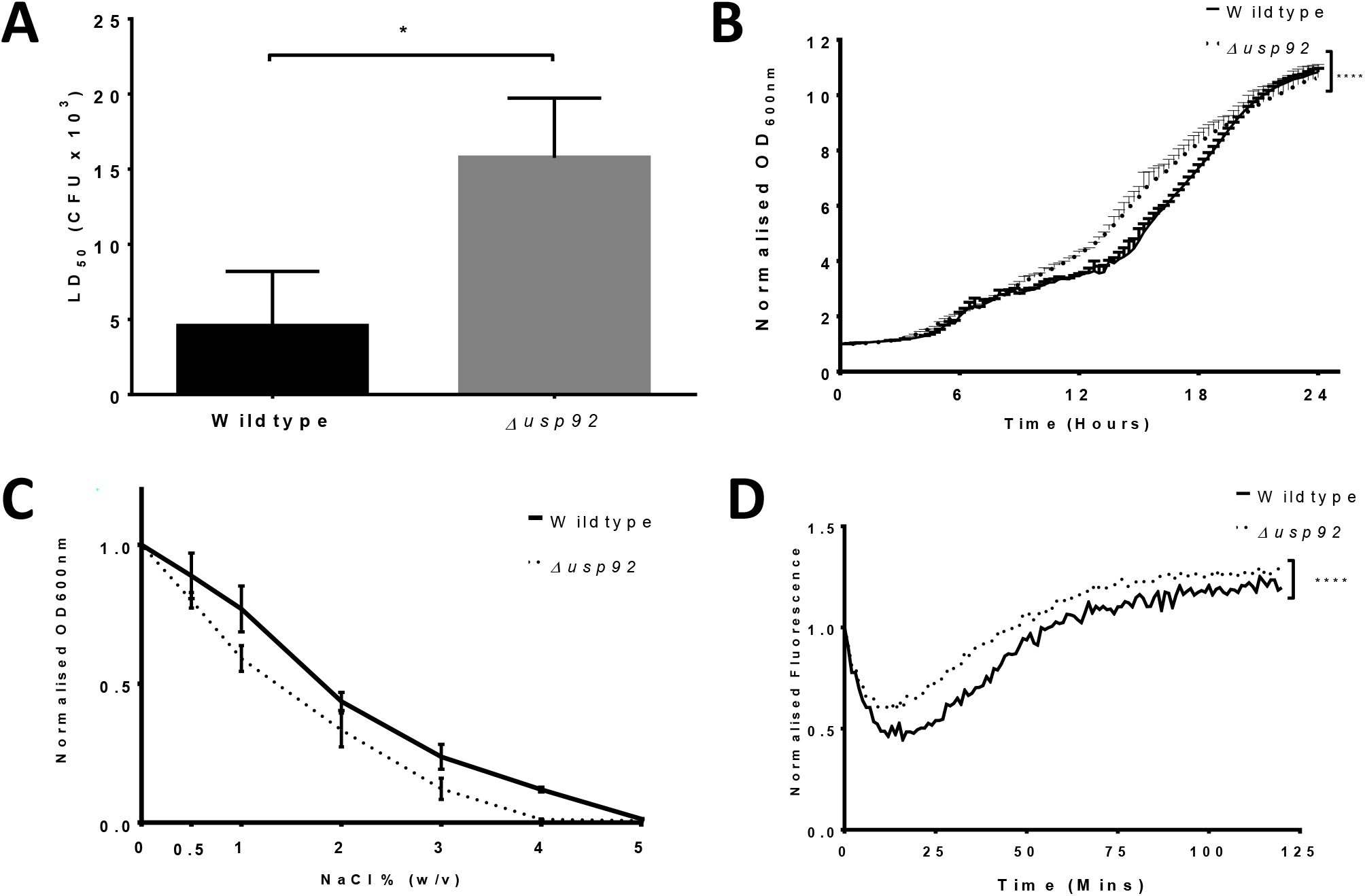
Phenotype analysis of USP92. A) Calculated LD_50_ values at 24 hours to determine virulence of the WT and the *Δusp92* mutant strains in the *G. mellonella* wax moth larva model Statistically significant difference relative to WT was determined by t-test, *p = 0.0228). **B)** Growth of WT and the *Δusp92* mutant following incubation with LB broth at a pH of 4.5 for 20 hours as determined by OD_600_. Data represent the mean OD values from three independent experiments at each timepoint. Error bars represent the standard error of the mean. Statistical significant difference determined by two-way ANOVA p < 0.0001. **C)** Mean endpoint absorbance values of WT and the *Δusp92* mutant following incubation with concentrations of NaCl (0 – 5% w/v) for 24 hours at 37°C normalised to LB only. *p=0.003 compared with wild type using two way ANOVA. **D)** Normalised fluorescence intensity data of the cellular uptake of Hoechst 33324 (excitation 355 nm, emission 460nm) in WT and the *Δusp92* mutant over a period of two hours, incubated at 37°C. Data represents the mean of eight replicates of each strain, performed on two independent experiments (*p < 0.001, as determined by two way ANOVA)

### Structural features of USP76 and USP92 explain their differential host cell attachment abilities

To identify structural determinants responsible for the different behaviours of USP76 and USP92, in particular their different abilities to bind to epithelial cells, we used homology modelling and analysed resulting structures. Both structures are highly reliable, given the high sequence identities with their template structures. The two protein structures share a similar dimeric organisation (Figure 8), consistent with their relatively high sequence similarity (sequence identity =41.8%). Each monomer is composed of an open-twisted, five-strand parallel β-sheet, sandwiched by two α-helices on each side of the sheet (α1-α4 in Figure 8) and present structural features of ATP binding proteins. In both models, a wide ATP binding cleft runs adjacent to the central β-sheet and contacts the α-helices α1, α3 and α4 of each monomer (Figure 8).

**Figure 8:**
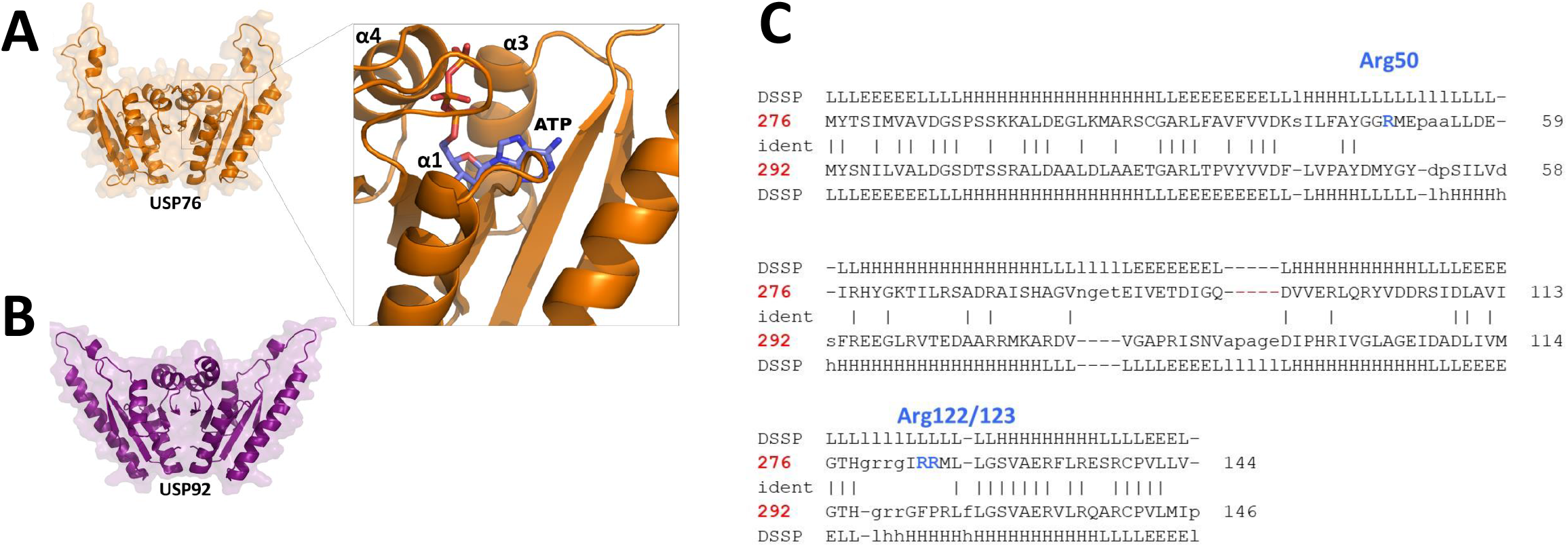
Comparison of USP76 and USP92 protein structures. a) Cartoon and surface representations of the homology models of (a) USP76 and (b) USP92. The inset shows a detail of ATP binding mode. (c) Structure based sequence alignment as computed by DALI.

Strong differences between the two proteins are evident when electrostatic potential surfaces are compared. Indeed, an overall negative electrostatic potential surface characterises USP92, with only two positively charged patches due to Arg119-120 and Arg136 (Figure 9). Consistent with a higher pI value of USP76 (pI=7.8) compared to USP92 (pI=5.0), the electrostatic potential surface of USP76 presents several clusters of positively charged residues on the entire surface of the protein, due mostly to arginine residues (Figure 9).

**Figure 9.**
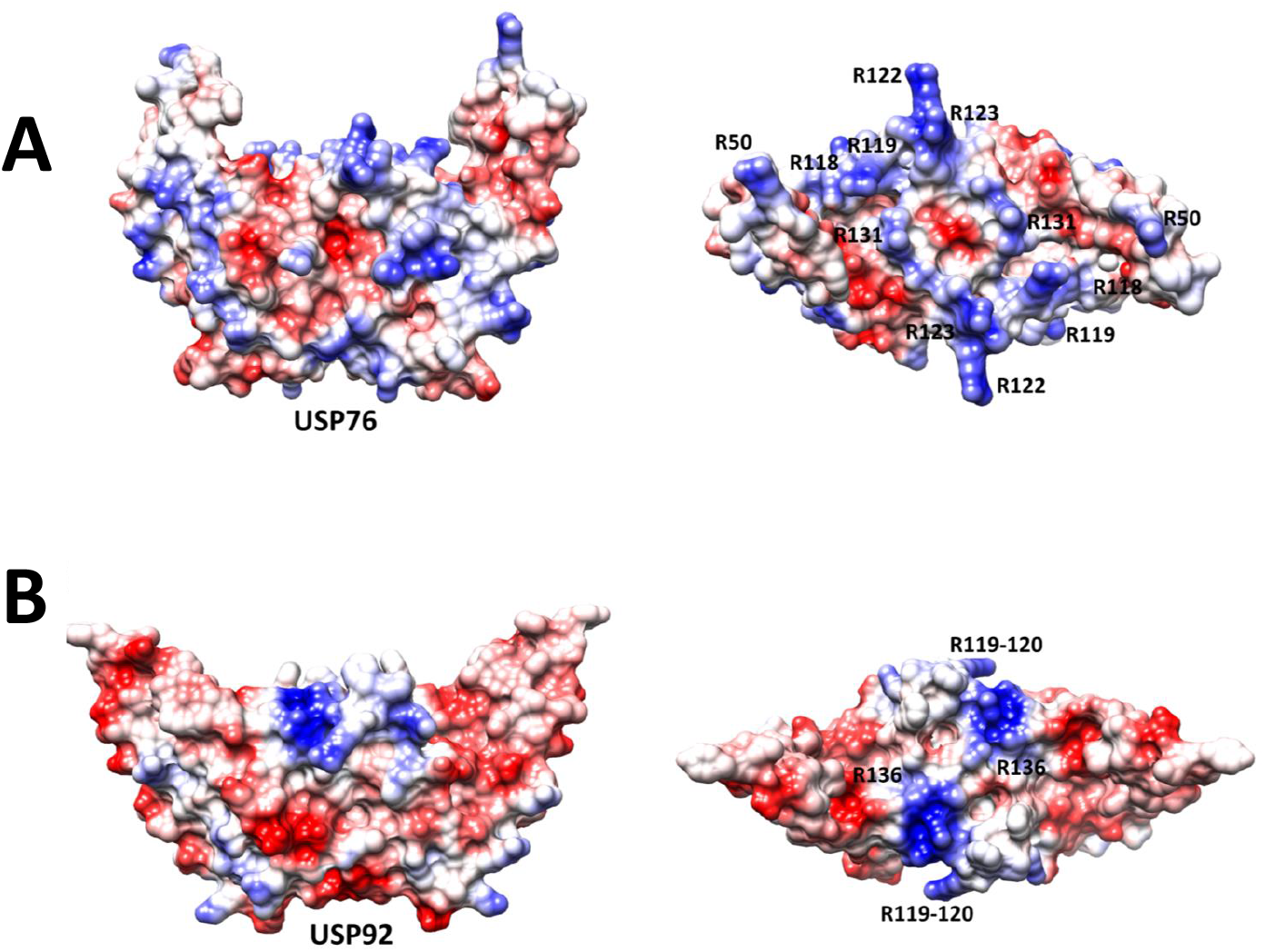
Side (left) and top (right) views of electrostatic potential surfaces of (a) USP76 and (b) USP92; with red and blue denoting residues with negative and positive electrostatic potential, respectively. The main residues contributing to the electrostatic potential are labelled.

## Discussion

A hallmark of genus Burkholderia is its remarkable ability to survive and thrive in a range of diverse environments, ranging from the soil, aquatic niches, disinfectants to the human host. Its ability to adapt to changing environments contributes to its success as a human pathogen. In this study we demonstrate that USPs support intramacrophage survival of *B. cenocepacia* and consequently may play an important role in the chronic colonisation of *B. cenocepacia* in people with cystic fibrosis. USPs are ubiquitous proteins which play a wide array of protective roles in bacterial pathogens and appear to be central to pathogenesis of intracellular pathogens. *B. cenocepacia* expresses 11 USPs. Six of these are encoded within the lxa locus, all of which increase in abundance in chronic infection and show increased expression in response to low oxygen [10, 11]. Despite the upregulation in response to low oxygen and chronic infection, we now show that USP76 and USP92 have quite distinct roles in *B. cenocepacia*, highlighting a clear lack of redundancy in these USPs. Moreover, we have shown that USP76, in particular, is likely to be important for survival of *B. cenocepacia* in CF macrophage, a significant characteristic of this pathogen.

To elucidate the role that both USPs play in Bcc chronic infection, we examined a series of phenotypes associated with environmental pressures experienced during chronic infection or associated with Bcc virulence. The CF lung has profoundly low pO2 due to a combination of disease associated issues including mucous plugging, constant neutrophilic inflammation and increased epithelial oxygen consumption [17, 26]. Oxidative stress in the CF lung contributes to the cycle of inflammation and is an inherent feature of CF [27]. Four key phenotypes relating to growth and/or survival under conditions typical of the CF lung were significantly altered in the *Δusp76* mutant compared to the wildtype strain: hypoxia, low pH, induced oxidative stress and nutrient starvation. Therefore, it is clear that USP76 (but not USP92) protects *B. cenocepacia* against oxidative stress. UspA gene deletion mutants in *Listeria monocytogenes* also had impaired growth when exposed to oxidative stress [28]. Similarly, *E. coli* UspA and UspD mutants were more susceptible to oxidative and superoxide stress [8, 29]. *L. monocytogenes* UspA mutants exposed to acidic stress were also previously found to have reduced cellular growth, albeit at a lower pH (pH 2.5) than that used by us [28]. In addition, an *A. baumannii* UspA mutant was also shown more susceptible to low pH and also oxidative stress [5]. Overexpression of a mycobacteria USP encoded by RV2624c increased survival in hypoxic conditions [30].

Previous reports showing that a *L. monocytogenes* UspA protected against low pH, oxidative stress and enhanced survival within murine macrophage [28], led us to evaluate whether USP76 also contributed to the survival of *B. cenocepacia* in macrophages, particularly given that USP76 was also protective in oxidative stress and involved in host cell attachment, in contrast to USP92. The impaired survival of the *Δusp76* mutant in U937 macrophage cells and, crucially, our finding that survival was significantly impaired in 80% of the CF-patient derived macrophage samples confirms that USP76 confers a clear survival advantage to *B. cenocepacia* inside macrophage cells. People with CF can have impaired macrophage function, which can lead to altered phagocytosis and killing of bacteria [31], consequently the role of USP76 in uptake and survival in CF-patient derived macrophages is particularly noteworthy. Given that impaired survival of the *Δusp76* was not evident in all CF-patient derived macrophages samples, it is clear that host factors also play a role. The BCAM0276 gene was previously shown to be upregulated when *B. cenocepacia* strain K56-2 was internalised in murine macrophages [32]. This work now confirms that USP76 contributes to survival of *B. cenocepacia* within the CF lung and also within the CF macrophage. Moreover, our previously observed increase in abundance of USP76 in chronically colonised patients indicates that USP76 is very likely to be a major player in facilitating the long-term survival of this pathogen within macrophages of CF patients.

The role of USP92 is distinct from that of USP76, as the two proteins are involved in determining different subsets of phenotypes (Table 2). Although both USPs protect the bacterial cell from acidic stress, only USP92 has a role in growth in the presence of osmotic stress. Moreover, we observe that USP92 is important for virulence in the acute larval infection model, in contrast to USP76, albeit not involved in host-cell attachment. The protective role against osmotic stress is critical in the context of CF airway surface liquid and may contribute to protection of *B. cenocepacia* in CF sputum. USPs in other species, e.g. atypical *E. coli*, have also been shown to be involved in salt tolerance [33].

**Table 2.**
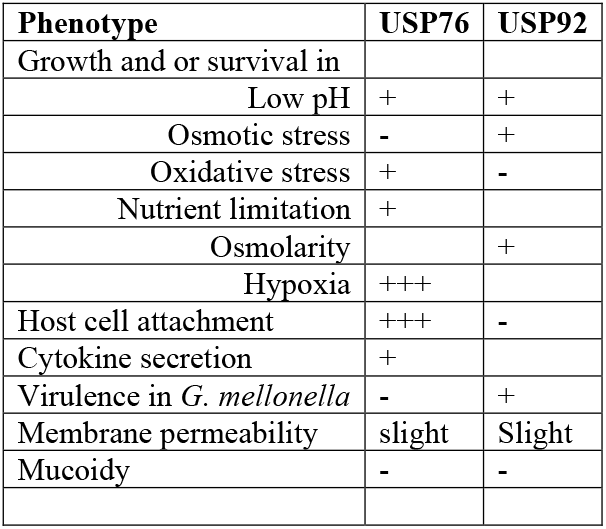
Comparison of phenotypes in USP76 and USP92.

Different roles for individual USPs expressed by a bacterial species have previously been observed in *E. coli*. Opposing roles in attachment have previously been reported for different USPs expressed in *E.coli*; with UspC and UspE mutants showing enhanced host cell attachment and loss of motility while UspF and UspG mutants had reduced host cell attachment and maintained cell motility [34]. Moreover, *E. coli* UspA and UspD were required for protection against oxidative stress while UspC and UspE were not. In *B. cenocepacia* opposing roles in membrane permeability were also observed for USP76 and USP92, with the *Δusp76* mutant having reduced permeability while the *Δusp92* mutant had increased permeability relative to the WT strain. A role in permeability would not be expected for cytosolic proteins that are expressed without signal peptides (likelihood of 0.0104) (SignalP-5.0). Yet the responses to Hoechst indicates that both these USPs may interact with the cellular membrane. The role of USP76 in host cell attachment strengthens this view. We have identified USPs, including USP76, among many cytosolic proteins in the outer membrane vesicles (OMVs) released from *B. cenocepacia* strain by proteomic analysis (unpublished data), and their presence in OMVs may provide a mechanism by which USPs can impact on the cell surface. As with USP76, *E.coli* USPs have also been linked to cell adhesion when mutants were assessed by yeast agglutination, with mutants of UspC and UspE enhancing cellular attachment, USPs F and G reducing in attachment [34]. The surface arginine residues on USP76 are likely to be involved in electrostatic interactions with sulfonate and carboxylate groups heparan sulphate, a negatively charged linear sulphated glycosaminoglycan (GAG) on the surface of epithelial cells. Indeed, it is well established that interactions of heparan sulphate with proteins are primarily driven by ion pair interactions mediated by surface regions rich in lysines and/or arginines, as in the case of the HBHA protein of *M. tuberculosis* [23–25]. Interestingly, macrophage cells increase expression of heparan sulphate under chronic inflammatory conditions, which would confer additional advantages on Bcc [36]. We speculate that USP76 is released from *B. cenocepacia* in OMVs, coating the bacterium and enhancing attachment to both macrophage and epithelial cells conferring a major advantage in its survival in the CF lung and increasing colonisation fitness. OMV released by *Vibrio cholerae* play a comparable role in surface adaptation in vivo [37], and this will hypothesis will need to be evaluated in *B. cenocepacia*.

USPs are clearly critical for an environmental bacterium such as *Burkholderia* to cope with the breadth of stressful conditions that it is exposed to in soil. However, it is now clear that USPs also confer substantial advantages on opportunistic pathogens adapting to the niche environment of the CF lung. In particular, USP76 is a major contributor to macrophage survival, while USP92 may confer advantages in the extracellular millieu, such as in CF sputum. Furthermore, the enhanced expression of both USPs over the course of chronic infection indicates that they play a role in the adaptation to chronic infection and represent interesting targets to overcome chronic colonisation.

## Experimental Procedures

### Bacterial Strains and Growth Conditions

Strains and plasmids used in this study are listed in table S1. Bacteria were routinely grown at 37^o^C in Luria-Broth (LB) medium with orbital shaking (200 rpm) unless otherwise stated. Antibiotics, when required, were added to reach final concentrations as follows: 50 μg/ml trimethoprim for *E. coli* and 100 μg/ml for *B. cenocepacia* and 40 μg/ml kanamycin for *E. coli*.

### Mammalian Cell Culture

Cystic fibrosis epithelial cells, CFBE41o^-^ which are homozygous for the ΔF508 mutation of the CFTR gene were routinely cultured in collagen/fibronectin coated flasks as previously described [12]. The U937 macrophage cells were maintained as a suspension culture in complete RPMI medium (Sigma-Aldrich) supplemented with 1 mM sodium pyruvate, 10 mM 4-(2-hydroxyethyl)-1-piperazineethanesulfonic acid (HEPES) buffer (pH 7.0 – 7.6), 1 mg /100 units streptomycin / penicillin, 10 % (v/v) FBS and 5 g/L D-glucose. U937 cells were plated in 24-well plates and after 24 h were induced to differentiate by the addition of 15 ng/ml of phorbol 12-myristate 13-acetate (PMA) for 24 hours in full RPMI medium.

### Construction of Δusp mutants and complementation

Targeted gene deletions of BCAM0276 or BCAM0292 in the *B. cenocepacia* strain K56-2 were performed as described [38]. The amplicons used to construct the mutagenic plasmid were cloned into pGPI-SceI-2 digested with *EcoRI* and *NheI*, using triparental mating, followed by biparental mating to introduce the I-SceI endonuclease. Screening was performed on the resulting colonies from bi-parental mating to determine if successful gene deletion had taken place. Trimethoprim sensitive colonies were screened for gene deletion using the US forward

DS reverse primers (Table S2) and mutants were sent for sequencing to confirm sequence deletion and stocks in glycerol prepared. To complement *B. cenocepacia K56-2ΔBCAM0276*, wild-type BCAM0276 was amplified from *B. cenocepacia* K56-2 with the complementation primer pair (Table S2), digested with the restriction enzymes *NdeI* and *XbaI* and ligated into similarly digested pMH447 [39]. The complementation plasmid was introduced into the mutant by conjugation. Once transferred into the target mutant strain the complementation vector integrates into the genome at aminoglycoside efflux genes (*BCAL1674-BCAL1675*), due to sequence homology between the vector and target genome [40]. As before, pDAI-Sce-I, was then introduced resulting in the replacement of BCAL1674-1675 by BCAM0276. The complementation of the BCAM0276 gene was confirmed by PCR and by phenotype analysis. Following genetic manipulation of *B. cenocepacia* K56-2 strain during gene deletion and subsequent complementation, the presence of the plasmid pC3 was confirmed using three sets of primers designed for genes RepA, oriC and dopC. The pC3 plasmid is a non-essential replicon of B. cenocepacia and encodes for a number of virulence factors [41], which can be lost during genetic manipulation. PCR was performed on confirmed mutants and complements using primers 3001/3002, 3003/3004 and 3005/3006.

### Bacterial attachment to human CF epithelial cells

CFBE41o^-^ cells were seeded in wells of a 24-well plate at a density of 4 x 10^5^ cells / well in antibiotic free medium and incubated overnight. The cells were then washed three times with PBS and the mid-logarithmic phase bacterial cultures (OD600nm 0.6-0.8) were resuspended in MEM and added to each well at a concentration of 2 x 10^7^ CFU / well (MOI 50:1). The plates were centrifuged at 252 x g for 5 min and incubated for 30 min at 37°C, 5% CO_2_ to allow for bacterial adherence. Wells were then washed with PBS and lysis buffer (0.25 % Triton X-100 in PBS) added to each well for 20 min at RT. Cell lysates were plated onto LB agar in duplicate and incubated at 37°C for 48 hours. Resulting colonies were counted and the CFU/ml determined. For microscopic visualisation of attachment, CFBE41o^-^ cells were seeded in chamber slides (LabTek™) and incubated with bacteria for 30 min and washed with PBS as outlined above. CFBE41o^-^ cells and adherent bacterial cells were fixed using 3 % w/v PFA (pH 7.2) for 10 min at RT, washed with PBS and blocked with 5 % BSA in PBS for one hour at RT. Cells were then incubated with a rabbit anti-Bcc antibody (courtesy of Prof U. Sajjan) in 1% BSA PBS overnight at 4°C. Cells were then washed with PBS and incubated with an anti-rabbit FITC conjugated antibody for 1 hour at 4°C. Cells were then washed twice with PBS for 5 min. Nuclei were counterstained with DAPI (VectaShield) and visualised using an Olympus FV-1000 confocal microscope. Bacterial attachment was expressed as number of bacteria per 100 cells in 10 randomly selected fields.

### Cytokine secretion by CFBE41o^-^ cells following infection with B. cenocepacia strains

CFBE41o^-^ cells were seeded at a density of 4 x 10^5^ cells / well in a 24-well plate in antibiotic medium for 24 hours followed by a further 24 hours in antibiotic and serum free medium prior to infection with *B. cenocepacia*. Overnight cultures of each strain were added to fresh LB medium and grown to an OD_600nm_ of between 0.4 – 0.8. Each strain (2 x 10^7^ CFU / ml, MOI 50:1) in MEM was added to each well in duplicate and incubated for 24 hours at 37°C and 5% CO_2_, then centrifuged at 315 x *g* for 12 min and supernatants transferred to −80°C for storage prior to assay. Interleukin-8 (IL-8) and IL-6 secretion was determined using OptEIA™ ELISA kits (Becton Dickenson) according to manufacturer’s instructions and the absorbance measured at 450 nm and 570 nm on the Biotek Synergy H1 Multiplate reader.

### Bacterial uptake and survival in U937 macrophage cells

The internalisation of *B. cenocepacia* strains by U937 macrophages was determined as previously described [42]. Briefly overnight cultures of each strain were transferred to fresh LB broth and incubated until mid-logarithmic phase (OD_600nm_ of 0.6 – 0.8) and diluted to an MOI of 5:1 (2.5 x 10^6^ CFU / ml) in RPMI medium. U937 cells were seeded at 5 x 10^5^ cells / ml in a 24-well plate, differentiated with PMA, washed twice with PBS, and 1 ml of each bacterial culture applied in duplicate. The CFU applied was confirmed by serial dilution of aliquots in Ringer’s and plating in duplicate. The 24-well plates were centrifuged at 1100 x _g_ for 5 min, incubated at 37°C, 5% CO_2_ for two hours, before washing with PBS. Extracellular bacteria were killed with amikacin/ceftazidime (1mg/ml each) for two hours at 37°C, 5% CO_2_. Wells were then washed 5 times with sterile PBS before cells were lysed with 0.25% Triton X-100 for 15 min and scraped, diluted and plated as outlined above. To examine intracellular survival of *B. cenocepacia* in U937 macrophage, bacteria were incubated with U937 as described above and after two hours incubation with each strain, the wells were washed PBS before addition of RPMI amikacin/ ceftazidime (1 mg/ml each) to each well and incubation for a further 2 hours at 37°C, 5% CO_2_.. The wells were subsequently washed twice with PBS and fresh antibiotics replaced and incubated at 37°C, 5% CO_2_ for various time points. Cells were then washed, lysed, diluted and plated as described above and CFU determined after 48 h. The % survival was calculated as the intracellular CFU/ml at each time point relative to the starting CFU/ml at time zero.

### Isolation of human peripheral mononuclear cells (PBMC) and differentiation

Ethical approval was obtained from the St Vincent’s University Hospital Research Ethics committee. Age and gender matched adults with CF with no history of Bcc infection were recruited from St Vincent’s University Hospital and blood samples were collected in EDTA tubes and diluted with the same volume of Dulbecco’s PBS (DPBS) and mixed by inversion.

PBMCs were isolated by layering the blood over using Ficoll-Paque plus (GE Healthcare) and centrifuging at 400 x *g* for 30 min at room temperature without braking. Upper layers containing plasma and platelets were removed and the layers containing mononuclear cells were transferred to a fresh tube containing three volumes of DPBS. Cells were centrifuged at 400 x *g* for 15 min at room temperature, resuspended in 8 ml DPBS and centrifuged at 100 x *g* for 10 min at to remove platelets. The cells were resuspended in RPMI, cryopreserved at 2 x 10^6^ cells per ml in 40% FBS, 10% DMSO, 50% RPMI (v/v) and stored in liquid nitrogen. Stored vials were revived as required and added to 5 ml RPMI 1640 (Sigma) medium (with no additives) and centrifuged at 100 x *g* for 15 min. The pellets were resuspended in 5 ml of RPMI 1640, transferred to a T25 and incubated at 37°C, 5% CO_2_ for two hours to allow the PBMCs to adhere to the flask. The cells were washed four times with warmed Mg^++^ and Ca^++^ free Dulbecco’s PBS. Full RPMI 1640 medium containing 10 % autologous Human Serum (Sigma), 10 mM HEPES (pH 7.0 – 7.6), 1mg / 100 units Penicillin / Streptomycin, 4.5 g/L D-glucose, 1 mM sodium pyruvate and 1 X non-essential amino acids (Sigma) were added to the flasks and the cells incubated with macrophage colony-stimulating factor (25 ng/ml, MCSF)(MSC) at 37°C, 5% CO_2_ for 7 days with half media changes every 3 days. The cells were harvested from the T25 flask by addition of porcine trypsin / EDTA (Sigma) in PBS and incubated at 37°C for 20 min. Cells were gently removed from the flask with a cell scraper and added to full RPMI medium and centrifuged at 252 x *g* for 10 min. Cells were resuspended in 1 ml of full RPMI medium, counted and diluted to the required concentration of cells in a 24-well plate. Plates were then incubated for 24 hours at 37°C prior use. Bacterial uptake and survival were then determined in the monocyte-derived macrophage cells as described for U937 cells.

### *Galleria mellonella* acute infection model

To determine the acute virulence of the bacterial strains, the *Galleria mellonella* wax moth larvae (Livefoods direct, Sheffield UK) were maintained at 15°C for seven days post-delivery prior to use and used within 4 weeks [43]. Overnight cultures of each bacterial strain were inoculated into 100 ml LB and grown to mid-logarithmic phase (OD_600nm_ of 0.6-0.8), diluted to ~ 1 x 10^6^ CFU / ml and centrifuged at 2500 x *g* for 10 min. Pellets were resuspended in 2 ml PBS and serially diluted to 10^-7^ in PBS and an aliquot of 20 μl of each dilution was injected into the hindmost left proleg of 10 healthy larvae weighing between 0.2 and 0.4 g using a sterile terumo 0.3 ml syringe. The bioburden injected was confirmed by serial dilution and plating. The larvae were then incubated at 37°C and the % survival of the larvae determined over 72h and plotted against the CFU bioburden inoculated value to calculate the LD_50_ value for each strain.

### Assessment of environmental stress responses

A number of environmental stresses experienced during chronic infection within the CF lung and macrophage environment were tested on each of the *B. cenocepacia* strains.

#### Controlled Hypoxia at 6% O_2_

LB was equilibrated in a controlled hypoxia chamber (Coy Laboratories) at 6% oxygen for 24h. Overnight cultures of the bacterial strains (10 ml) were centrifuged at 4000 x *g* for 15 min, resuspended in 10 ml fresh LB, transferred to 40 ml hypoxia equilibrated LB in 100 ml conical flask, and incubated statically at 37°C and 6 % O_2_. Aliquots were sampled at 24 hour intervals over 8 days, serially diluted to 10^-7^ in Ringer’s solution, plated onto LB agar plates in duplicate and then incubated at 37°C for 48 hours in normoxic conditions before enumeration.

#### Low pH

Overnight cultures were diluted 1:100 LB broth at either pH 4.5 or standard pH ~7.5 and 300 μl aliquots added in duplicate to the wells of a 96-well round bottomed plate. The plates were incubated at 37°C, with orbital shaking and OD_600nm_ measured every 15 min for 24 hours. Cell viability in low pH was also determined by inoculating a 100 ml flask of LB broth at pH 4.5 with 10 ml of an overnight culture and incubated at 37°C, 170 rpm and plating hourly samples as described previously.

#### High osmolarity

The effect of high osmolarity was determined in each *B. cenocepacia* strain by adding 300 μl of a range of concentrations of NaCl (0 – 5% w/v) or sucrose (0 – 50%) to rows of a 96-well plate. Overnight cultures were added to each well (3 μl) in duplicate and the plates were incubated in a Biotek Synergy H1 multiplate reader at 37°C for 24 hours with OD_600nm_ measured every 15 min.

#### Oxidative Stress

The effects of oxidative stress on the *B. cenocepacia* strains was assessed firstly by exposing the strains to a series of concentrations of organic peroxide (*tert*-butyl hydroperoxide) and inorganic peroxide (hydrogen peroxide, H_2_O_2_). Overnight cultures of each strain were diluted 1:100 in fresh LB broth and 270 μl added in duplicate to corresponding wells in a 96-well plate. A series of concentrations of either H_2_O_2_ (0 – 1 mM) or *tert*-butyl hydroperoxide (0 – 200 μM) were added to the wells and growth determined in a Syngery H1 microplate reader at 37°C at medium shaking and OD_600nm_ measured every 15 min for 20 hours. Viability of bacterial cells in an oxidative stress environment was also examined by treating cultures with 700 μM H_2_O_2_ and incubated at 37°C, 200 rpm. Hourly samples were serially diluted in Ringer’s solution, and plated in duplicate to determine CFU / ml.

#### Heat Stress at 42°C

The effect of heat on each *B. cenocepacia* strain was determined by transferring 10 ml of overnight cultures of each strain into 100 ml of pre-warmed (42°C) LB and incubation at 42°C, 200 rpm. Hourly samples were diluted in Ringer’s solution, plated and enumerated after 48h.

#### Long term Nutrient Starvation

Long term nutrient starvation on each *B. cenocepacia* strain was assessed as per Silva *et al* (2013). Overnight cultures of each strain grown in LB broth were centrifuged at 4000 x *g* for 5 min and pellets were washed twice in 0.9 % w/v NaCl and resuspended in 9 ml of M63 medium (2 g/L ammonium sulphate, 13.6 g/L potassium phosphate monobasic, 0.5 mg/L Iron (II) sulphate, 0.2 g/L Magnesium sulfate) with no carbon source. This was then added to 41 ml M63 medium in a 100 ml conical flask and incubated at 37°C with agitation of 200 rpm. Aliquots were taken for 28 days, serially diluted in Ringer’s solution, plated and counted after 48h incubation.

#### Assessment of cell permeability by Hoechst 33324

The cellular permeability of each strain was assessed by diluting overnight cultures 1:10 in fresh LB medium, incubating at 37°C, 200 rpm for 5 hours, centrifuging at 4,000 x *g* for 3 min and resuspending in sterile PBS. Each strain was then diluted to an OD_600nm_ of 0.1 with sterile PBS and 180 *μ*l added to 8 replicate wells of a row in a black, fluorescence 96-well plate and 2.5 μM of Hoechst 33324 added [44]. The plates were incubated in a Biotek Synergy H1 Multiplate reader at 37°C, medium shaking and fluorescence was measured every minute for 2 hours at an excitation of 355 nm and an emission of 460 nm. The mean fluorescence of 8 replicate wells was calculated for each strain and normalised with the T_o_ value.

### Homology modelling

The homology model structures of BCAM0276 and BCAM0292 were obtained after consensus-based sequence alignment using the HHpred tool. The best model template for BCAM0276 was identified as the structure of the TeaD stress protein from the TRAP transporter TeaABC of *Halomonas elongata* (PDB code 3hgm, seqid 34.3%). For BCAM0292, the best template was the BupsA stress protein from *Burkholderia pseudomallei* (PDB code 4wny, seqid 69.2%). Using these alignments, the homology models were built using the program MODELLER [45]. Electrostatic potential surfaces were computed using the program Chimera [46].

### Statistical analysis

Statistical analysis of host cell attachment, cytokine secretion, CF macrophage uptake and survival data was performed by one-way ANOVA using Prism software. Two-way ANOVA was used to analyze growth in the presence of organic and inorganic peroxide, growth under acidic conditions and nutrient limitation. Statistical analysis of virulence, survival at 8 days in hypoxic conditions, endpoint OD_600_ in acidic conditions, U937 internalisation were performed using a student’s t-test.

Assessment of antibiotic resistance, mucoidy, motility, biofilm formation was performed as described in the Supplementary information.

## Acknowledgements

We are most grateful to Prof Umadevi Sajjan (University of Michigan) for the gift of the anti-Bcc antibody. This research was supported by a President’s Award from the Institute of Technology Tallaght, Dublin and a Government of Ireland Postgraduate scholarship Award (GOIPG/2017/788) from the Irish Research Council and EU COST Action BM1003: “Microbial cell surface determinants of virulence as targets for new therapeutics in cystic fibrosis for Short Term Scientific Missions”.

## References

1. O’Connor A, McClean S. The Role of Universal Stress Proteins in Bacterial Infections. Curr Med Chem 2017; 24:3970–9.

2. Schreiber K, Boes N, Eschbach M, et al. Anaerobic survival of Pseudomonas aeruginosa by pyruvate fermentation requires an Usp-type stress protein. J Bacteriol 2006; 188:659–68.

3. Boes N, Schreiber K, Härtig E, Jaensch L, Schobert M. The Pseudomonas aeruginosa universal stress protein PA4352 is essential for surviving anaerobic energy stress. J Bacteriol 2006; 188:6529–38.

4. Drumm JE, Mi K, Bilder P, et al. Mycobacterium tuberculosis universal stress protein Rv2623 regulates bacillary growth by ATP-Binding: requirement for establishing chronic persistent infection. PLoS pathogens 2009; 5:e1000460.

5. Elhosseiny NM, Amin MA, Yassin AS, Attia AS. Acinetobacter baumannii universal stress protein A plays a pivotal role in stress response and is essential for pneumonia and sepsis pathogenesis. Int J Med Microbiol 2015; 305:114–23.

6. Nyström T, Neidhardt FC. Effects of overproducing the universal stress protein, UspA, in Escherichia coli K-12. J Bacteriol 1996; 178:927–30.

7. Nyström T, Neidhardt FC. Cloning, mapping and nucleotide sequencing of a gene encoding a universal stress protein in Escherichia coli. Molecular microbiology 1992; 6:3187–98.

8. Nyström T, Neidhardt FC. Expression and role of the universal stress protein, UspA, of Escherichia coli during growth arrest. Molecular microbiology 1994; 11:537–44.

9. Vanlaere E, Baldwin A, Gevers D, et al. Taxon K, a complex within the Burkholderia cepacia complex, comprises at least two novel species, Burkholderia contaminans sp. nov. and Burkholderia lata sp. nov. Int J Syst Evol Microbiol 2009; 59:102–11.

10. Sass AM, Schmerk C, Agnoli K, et al. The unexpected discovery of a novel low-oxygen-activated locus for the anoxic persistence of Burkholderia cenocepacia. Isme J 2013; 7:1568–81.

11. Cullen L, O’Connor A, McCormack S, et al. The involvement of the low-oxygen-activated locus of Burkholderia cenocepacia in adaptation during cystic fibrosis infection. Scientific Reports 2018; 8:13386.

12. Cullen L, O’Connor A, Drevinek P, Schaffer K, McClean S. Sequential Burkholderia cenocepacia Isolates from Siblings with Cystic Fibrosis Show Increased Lung Cell Attachment. American journal of respiratory and critical care medicine 2017; 195:832–5.

13. Nunvar J, Kalferstova L, Bloodworth RA, et al. Understanding the Pathogenicity of Burkholderia contaminans, an Emerging Pathogen in Cystic Fibrosis. PLoS One 2016; 11:e0160975.

14. Winsor GL, Khaira B, Van Rossum T, Lo R, Whiteside MD, Brinkman FS. The Burkholderia Genome Database: facilitating flexible queries and comparative analyses. Bioinformatics 2008; 24:2803–4.

15. Elizur A, Cannon CL, Ferkol TW. Airway inflammation in cystic fibrosis. Chest 2008; 133:489–95.

16. Roesch EA, Nichols DP, Chmiel JF. Inflammation in cystic fibrosis: An update. Pediatr Pulmonol 2018; 53:S30–S50.

17. Montgomery ST, Mall MA, Kicic A, Stick SM, Arest CF. Hypoxia and sterile inflammation in cystic fibrosis airways: mechanisms and potential therapies. Eur Respir J 2017; 49.

18. Martin DW, Mohr CD. Invasion and intracellular survival of Burkholderia cepacia. Infection and immunity 2000; 68:24–9.

19. Rosales-Reyes R, Aubert DF, Tolman JS, Amer AO, Valvano MA. Burkholderia cenocepacia type VI secretion system mediates escape of type II secreted proteins into the cytoplasm of infected macrophages. PLoS One 2012; 7:e41726.

20. Sousa MC, McKay DB. Structure of the universal stress protein of Haemophilus influenzae. Structure 2001; 9:1135–41.

21. Grandjean Lapierre S, Phelippeau M, Hakimi C, et al. Cystic fibrosis respiratory tract salt concentration: An Exploratory Cohort Study. Medicine (Baltimore) 2017; 96:e8423.

22. Zlosnik JE, Hird TJ, Fraenkel MC, Moreira LM, Henry DA, Speert DP. Differential mucoid exopolysaccharide production by members of the Burkholderia cepacia complex. J Clin Microbiol 2008; 46:1470–3.

23. Esposito C, Cantisani M, D’Auria G, et al. Mapping key interactions in the dimerization process of HBHA from Mycobacterium tuberculosis, insights into bacterial agglutination. FEBS Lett 2012; 586:659–67.

24. Esposito C, Marasco D, Delogu G, Pedone E, Berisio R. Heparin-binding hemagglutinin HBHA from Mycobacterium tuberculosis affects actin polymerisation. Biochem Biophys Res Commun 2011; 410:339–44.

25. Joseph PRB, Sawant KV, Iwahara J, Garofalo RP, Desai UR, Rajarathnam K. Lysines and Arginines play non-redundant roles in mediating chemokine-glycosaminoglycan interactions. Sci Rep 2018; 8:12289.

26. Mendelsohn L, Wijers C, Gupta R, Marozkina N, Li C, Gaston B. A novel, noninvasive assay shows that distal airway oxygen tension is low in cystic fibrosis, but not in primary ciliary dyskinesia. Pediatr Pulmonol 2019; 54:27–32.

27. Scholte BJ, Horati H, Veltman M, et al. Oxidative stress and abnormal bioactive lipids in early cystic fibrosis lung disease. J Cyst Fibros 2019; 18:781–9.

28. Seifart Gomes C, Izar B, Pazan F, et al. Universal stress proteins are important for oxidative and acid stress resistance and growth of Listeria monocytogenes EGD-e in vitro and in vivo. PLoS One 2011; 6:e24965.

29. Nachin L, Nannmark U, Nyström T. Differential roles of the universal stress proteins of Escherichia coli in oxidative stress resistance, adhesion, and motility. J Bacteriol 2005; 187:6265–72.

30. Jia Q, Hu X, Shi D, et al. Universal stress protein Rv2624c alters abundance of arginine and enhances intracellular survival by ATP binding in mycobacteria. Sci Rep 2016; 6:35462.

31. Leveque M, Le Trionnaire S, Del Porto P, Martin-Chouly C. The impact of impaired macrophage functions in cystic fibrosis disease progression. J Cyst Fibros 2017; 16:443–53.

32. Tolman JS, Valvano MA. Global changes in gene expression by the opportunistic pathogen Burkholderia cenocepacia in response to internalization by murine macrophages. BMC Genomics 2012; 13:63.

33. de Souza CS, Torres AG, Caravelli A, Silva A, Polatto JM, Piazza RM. Characterization of the universal stress protein F from atypical enteropathogenic Escherichia coli and its prevalence in Enterobacteriaceae. Protein Sci 2016; 25:2142–51.

34. Nachin L, Nannmark U, Nystrom T. Differential roles of the universal stress proteins of Escherichia coli in oxidative stress resistance, adhesion, and motility. J Bacteriol 2005; 187:6265–72.

35. Huang TY, Irene D, Zulueta MM, et al. Structure of the Complex between a Heparan Sulfate Octasaccharide and Mycobacterial Heparin-Binding Hemagglutinin. Angew Chem Int Ed Engl 2017; 56:4192–6.

36. Swart M, Troeberg L. Effect of Polarization and Chronic Inflammation on Macrophage Expression of Heparan Sulfate Proteoglycans and Biosynthesis Enzymes. J Histochem Cytochem 2018:22155418798770.

37. Zingl FG, Kohl P, Cakar F, et al. Outer Membrane Vesiculation Facilitates Surface Exchange and In Vivo Adaptation of Vibrio cholerae. Cell Host Microbe 2020; 27:225–37 e8.

38. Flannagan RS, Linn T, Valvano MA. A system for the construction of targeted unmarked gene deletions in the genus Burkholderia. Environ Microbiol 2008; 10:1652–60.

39. Hamad MA, Di Lorenzo F, Molinaro A, Valvano MA. Aminoarabinose is essential for lipopolysaccharide export and intrinsic antimicrobial peptide resistance in Burkholderia cenocepacia(dagger). Molecular microbiology 2012; 85:962–74.

40. Hamad MA, Skeldon AM, Valvano MA. Construction of aminoglycoside-sensitive Burkholderia cenocepacia strains for use in studies of intracellular bacteria with the gentamicin protection assay. Appl Environ Microbiol 2010; 76:3170–6.

41. Agnoli K, Frauenknecht C, Freitag R, et al. The third replicon of members of the Burkholderia cepacia Complex, plasmid pC3, plays a role in stress tolerance. Appl Environ Microbiol 2014; 80:1340–8.

42. McKeon S, McClean S, Callaghan M. Macrophage responses to CF pathogens: JNK MAP kinase signaling by Burkholderia cepacia complex lipopolysaccharide. FEMS Immunol Med Microbiol 2010:DOI:10.1111/j.574-695X.2010.00712.x.

43. Costello A, Herbert G, Fabunmi L, et al. Virulence of an emerging respiratory pathogen, genus Pandoraea, in vivo and its interactions with lung epithelial cells. J Med Microbiol 2011; 60:289–99.

44. Coldham NG, Webber M, Woodward MJ, Piddock LJ. A 96-well plate fluorescence assay for assessment of cellular permeability and active efflux in Salmonella enterica serovar Typhimurium and Escherichia coli. J Antimicrob Chemother 2010; 65:1655–63.

45. Bitencourt-Ferreira G, de Azevedo WF, Jr. Homology Modeling of Protein Targets with MODELLER. Methods Mol Biol 2019; 2053:231–49.

46. Yang Z, Lasker K, Schneidman-Duhovny D, et al. UCSF Chimera, MODELLER, and IMP: an integrated modeling system. J Struct Biol 2012; 179:269–78.

