## Supplemental data for "A Universal Stress Protein upregulated by hypoxia may contribute to chronic lung colonisation and intramacrophage survival in cystic fibrosis"

### Supplementary information:

**Supplementary materials and methods:**

##### Assessment of Antibiotic Susceptibility

Minimum inhibitory concentrations (MIC) were determined using Ceftazidime (0.016 – 256 μg), Levofloxacin (0.002 – 32 μg), Meropenem (0.002 – 32 μg), Polymyxin B (0.064 – 1024 μg) ETEST® strips (Biomerieux). Overnight cultures were diluted to $OD_{600nm}$ of 0.1 in sterile PBS and 100 μl aliquots spotted onto individual Muellar Hinton (MH) agar plates and using a PBS-wetted sterile cotton swab, the cultures were then spread horizontally, vertically and along the edge of the MH agar plates. Each ETEST® was placed in the centre of the individual MH agar plates and incubated at 37°C for up to 48 hours. The MIC value for each strain was determined by examining the elliptic inhibition zone intersecting the MIC value scale. The susceptibility of the strains to Polymyxin B was further examined by diluting overnight cultures 1:100 in LB broth and adding 270 μl to each well in duplicate in a 96-well plate. A series of Polymyxin B concentrations, prepared in distilled water with 0.2% (w/v) BSA and 0.02% (v/v) acetic acid were serially diluted 1:10 to give a final concentration in the range of 16 to 1024 μg/ml and the plate incubated in a Biotek Synergy H1 Microplate reader at 37°C with medium shaking and $OD_{600nm}$ measured every 15 min for 20 hours. [1]

##### Assessment of Mucoid Phenotype

Exopolysaccharide (EPS) production was assessed to determine mucoid phenotype of the mutant strains of Bcc using Yeast extract mannitol (YEM) (2g/L Yeast Extract, 20g/L D-Mannitol, 15g/L Agar no.2) plates [2]. Overnight cultures of each strain were streaked onto YEM plates in a quadrant streak manner and incubated at 37°C for 5 days in normal oxygen conditions or low oxygen conditions using a controlled hypoxic chamber (COY). EPS production and mucoid phenotype were scored as follows: The scoring of mucoid phenotype was as follows: (-) bacterial colonies on YEM plates appeared dry with a matte appearance showed no evidence of EPS production, (+) bacterial colonies predominately non-mucoid with some evidence of EPS production in the confluent area of growth on the YEM plate, (++) mucoid phenotypes with EPS production on both the confluent area of growth and streaked individual colonies which appeared flat on the YEM plates, (+++) excessive EPS production with poor distinction between confluent area and streaked area, streaks appeared raised due to EPS production; a score of (+++d) was similar to (+++) but the EPS production resulted in drips of EPS on the lid of the petri dish when inverted [3].

**Determination of motility**

Motility assays were performed on each mutant strain to determine the effect of gene deletion on the ability of swimming, swarming and twitching [4]. All agar plates were poured on a flat, even surface and allowed to dry for 30 min aseptically under laminar flow. Swimming motility was assessed by inoculating the surface of the agar plate (0.3% w/v LB agar) with 5 μl of an overnight culture using a sterile yellow 20 μl pipette tip. Plates were wrapped in parafilm, inverted and incubated at 37°C for 48 hours after which the average of two diameter measurements of the zone motility at 24 and 48 hours were measured. Strains were scored as follows: non-motile (diameter ≤ 5 mm), motile (diameter > 5 mm and ≤ 60 mm) or highly motile (diameter of >60 mm). Swarming motility was assessed by piercing 5 μl of an overnight culture into the centre of the (0.5% w/v LB agar) with a sterile yellow 20 μl pipette tip without piercing all the way through the agar. Plates were wrapped in parafilm, not inverted and incubated at 30°C for 48 hours and the average of two diameter measurements of the zone motility at 24 and 48 hours determined. Strains were scored as follows: non-motile (diameter ≤ 5 mm), motile (diameter > 5 mm and ≤ 60 mm) or highly motile (diameter of >60 mm). Twitching motility was assessed by inoculating the agar (1% w/v LB agar) with 5 μl of an overnight culture as above and incubating at 37°C for 24 hours. Plates were removed from the incubator and dried under laminar flow for 30 min, the agar removed before staining the plates with 0.5% w/v Coomassie brilliant blue R250, 40 % v/v Methanol and 10 % v/v Acetic Acid for 2 min. Excess stain was then washed off with a solution of 40 % methanol and 10 % acetic acid. Twitching was determined by taking the average of two diameter measurements of the zone motility. Strains were scored as follows: non-motile (diameter ≤ 5 mm), motile (diameter > 5 mm and ≤ 30 mm) or highly motile (diameter of >30 mm).

**Assessment of Biofilm formation**

Overnight cultures (10 ml) of each strain were transferred to 100 ml of fresh LB broth and incubated at 37°C for 4 – 5 hours with agitation at 200 rpm. $OD_{600nm}$ was measured and cells were diluted to ~

1.6 x $10^6$ CFU / ml in LB broth. Each strain (200 μl) was added to each of the 8 wells in the respective columns of triplicate 96-well round bottom plate with sterile LB broth used as the negative control. Plates were incubated at 37°C in normoxic or hypoxic (6 % $O_2$) conditions for up to 72 hours. Plates were removed at 24, 48 and 72 h intervals, wells washed twice with sterile water before addition of 125 μl of 0.1 % w/v crystal violet solution to stain biofilm mass for 30 min. The wells were washed 3 times with sterile water and allowed to dry for 30 min at RT, before addition of 100 μl of 95 % v/v ethanol to solubilise the stained biofilm for 15 min. The solution was transferred to a fresh 96-well flat bottom plate and the $OD_{590nm}$ was measured. Wells with an OD reading of ≤0.120 are considered non-biofilm forming. OD readings of ≥0.120 but <0.240 are considered poor biofilm formation and OD readings of >0.240 are strong biofilm formation.

Table S1. Bacterial strains and plasmids used in this study:

| | **Characteristics** | **Source** |
|---|---|---|
| **Strain** | | |
| ***B. cenocepacia*** | | |
| K56-2 | ET12 Clone related to J2315 | CF Clinical isolate, Canada (Epidemic strain) |
| K56-2 *Δusp76* | Deletion of BCAM0276 in K56-2 | This study |
| *K56-2 Δusp76_usp76* | BCAM0276 integration in K56-2 Δ0276, $Gn^S$ | This study |
| K56-2 *Δusp92* | Deletion of BCAM0292 in K56-2 | This study |
| **Plasmids** | | |
| pGPI-SceI-2 | *ori*R6K, $\Omega Tp^r$ , $mob^+$, containing the ISce-I restriction site | (Flannagan *et al.,* 2008) |
| pRK2013 | *ori*COIEI, RK2 derivative, $Kan^r$, $mob^+$, $tra^+$ | (Figurski and Helinski, 1979) |
| pDAI-SceI | Encodes the ISce-I homing endonuclease,*ori*pBBR1, $Tet^R$, Pdhfr, *$mob^+$* | (Flannagan *et al.,* 2008) |
| pMH447 | pGPI-SceI derivative used for chromosomal complementation, $Tp^R$ | This study (as per Hamad *et al.*, 2012) |

Table S2: The primers used in this study, underlined sequences indicate restriction sites.

| Primer No. | Primer Description | 5’ to 3’ Primer Sequence | Restriction Enzyme |
|---|---|---|---|
| 2761 | BCAM0276 Upstream | TTGAGAATTCAAAGTCACTCGGCTACCC | *ApaI* |
| 2762 | | AATTATCGATGAACAGCCGTGCGCC | *HindIII* |
| 2763 | BCAM0276 Downstream | TTTAATCGATGCTCCTCGGCAGTGTCG | *HindIII* |
| 2764 | | AAAAGCTAGCGCGGCCGATCACGTACAGAA | *NheI* |
| 2765 | BCAM0276 Internal Gene | CAGTAGACGGTAGTCCTTCATCAAAAA | |
| 2766 | | GATATCCGTCTCGACAATTTCCGTCTC | |
| 2767 | BCAM0276 Complementation | TTTTCATATGGAAACACATTCGAACGCC | *NdeI* |
| 2768 | | GAAATCTAGATCCGGCCCGCACGTTC | *XbaI* |
| 2921 | BCAM0292 Upstream | TCCGGAATTCTGTCGGGTGACGCGCT | *EcoRI* |
| 2922 | | GCTATAAGCTTCCGTCGAGCGCAACC | *HindIII* |
| 2923 | BCAM0292 Downstream | CAAGAAGCTTCGAGGCCACGGCTGC | *HindIII* |
| 2924 | | GGCAGCTAGCCGTCCTCGACGAAAT | *NheI* |
| 2925 | BCAM0292 Internal Gene | AAGGATTACGCGTGACCGAG | |
| 2926 | | ACGGTTCCTTTTCGGTGGAT | |
| 2927 | BCAM0292 Complementation | TCTGCATATGATGTACTCGAACATT | *NdeI* |
| 2928 | | TTAATCTAGATCATGACGGTTCCTT | *XbaI* |
| 4019 | pGPI-SceI / Sequencing | GAGTGACACAGGAACACT | |
| 4021 | | GCTCAATCAATCACCGGATCCC | |
| 5386 | pMH447 Construction | GGATCGAATTCCGGCTCGCCGCATTCCTGCAGG | *EcoRI* |
| 5387 | | TCCGATCCATATGGTTCTGACCTCGGATCATTCG | *NdeI* |
| 5388 | | CTCGCCAGCTAGCGCCGATATAGTCGGAGCCGAAC ATC | *NheI* |
| 5389 | | TAGTGCTCATATGGTCGTTTCTAGATGCTCACGCAA CTGCCGACCGC | *NdeI / XbaI* |
| 5885 | pMH447 Sequencing | TTGATGGCGAGCGATTCTTC | |
| 5886 | | CCAGTTCTTCAGCGTGACGA | |
| 3001 | Confirmation of PC3 plasmid | AACAGCAACAACACCAAGTA | |
| 3002 | | TTGCTTGCCTTCTTCTTCG | |
| 3003 | | ACCCATTTATGGGAGCACAG | |
| 3004 | | TGCTCCAGCATCACTTTCAC | |
| 3005 | | CGGGCTGAACAGTAACAATA | |
| 3006 | | GCTTGCTCGCTTTCTTTTTC | |

**Table S3:** EPS production score of the WT and the $\Delta$*usp76* mutant assessed on YEM agar plates. EPS production score of the WT and the $\Delta$*usp76* mutantstrains assessed following incubation on YEM agar plates for 5 days at 37°C on 3 independent occasions. EPS scored according to method described by Zlosnik *et al.,* 2008.

| **Strain** | **EPS Score (-/+/++/+++/+++d)** | **Description** |
| --- | --- | --- |
| WT | + | Some evidence of EPS |
| $\Delta$*usp76* | + | Some evidence of EPS |
| WT at 6% $O_2$ | + | Some evidence of EPS |
| $\Delta$*usp76* at 6% $O_2$ | + | Some evidence of EPS |

1. Loutet SA, El-Halfawy OM, Jassem AN, et al. Identification of synergists that potentiate the action of polymyxin B against Burkholderia cenocepacia. Int J Antimicrob Agents **2015**; 46:376-80.
2. Lagatolla C, Skerlavaj S, Dolzani L, et al. Microbiological characterisation of Burkholderia cepacia isolates from cystic fibrosis patients: investigation of the exopolysaccharides produced. FEMS Microbiol Lett **2002**; 209:99-106.
3. Zlosnik JE, Hird TJ, Fraenkel MC, Moreira LM, Henry DA, Speert DP. Differential mucoid exopolysaccharide production by members of the Burkholderia cepacia complex. J Clin Microbiol **2008**; 46:1470-3.
4. Cullen L, Weiser R, Olszak T, et al. Phenotypic characterization of an international Pseudomonas aeruginosa reference panel: strains of cystic fibrosis (CF) origin show less in vivo virulence than non-CF strains. Microbiology **2015**; 161:1961-77.

### Supplemental figs

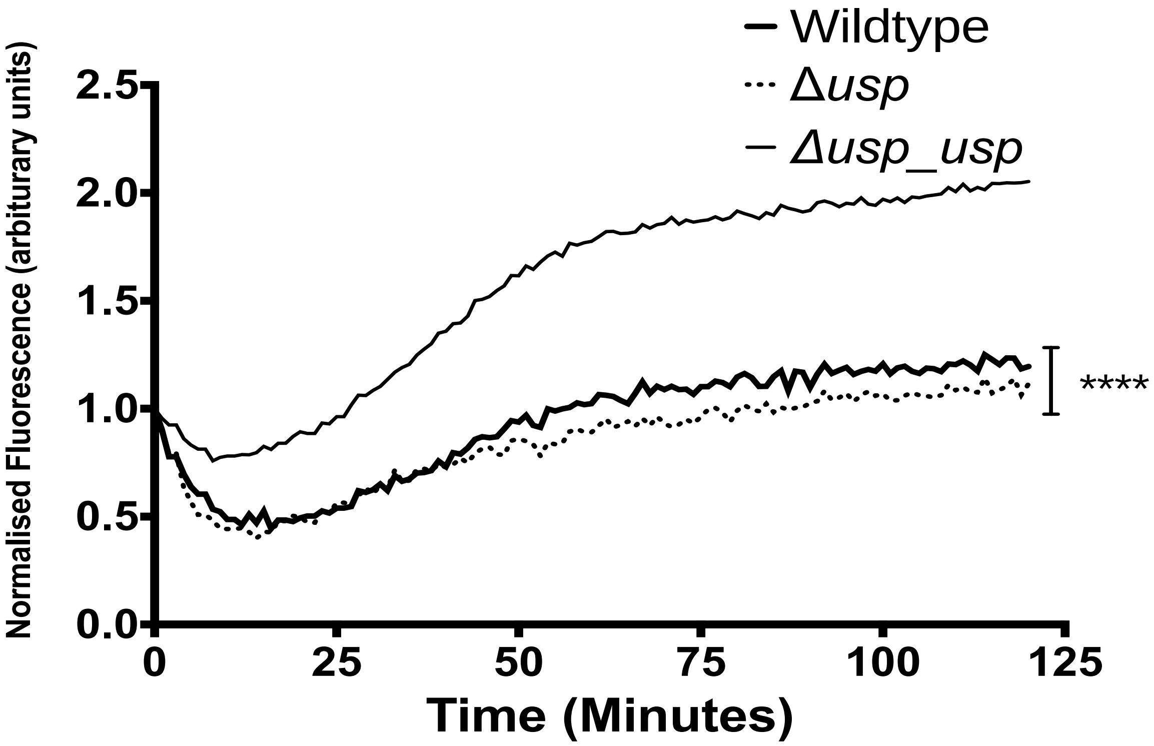

**Fig S1: Normalised fluorescence intensity for the measurement of cell permeability of the WT, the Δ*usp76* mutant and the Δ*usp76_usp76* complement strains with Hoescht 33324.** Normalised fluorescence intensity data of the cellular uptake of Hoescht 33324 (excitation 355 nm, emission 460nm) in WT, the Δ*usp76* mutant and the Δ*usp76_usp76* complement over a period of two hours, incubated at 37 °C. Data represents the average of eight replicates of each strain, performed on two independent occasions. **** Statistically significant difference relative to the WT as determined by two-way ANOVA ($p = 0.0001$)

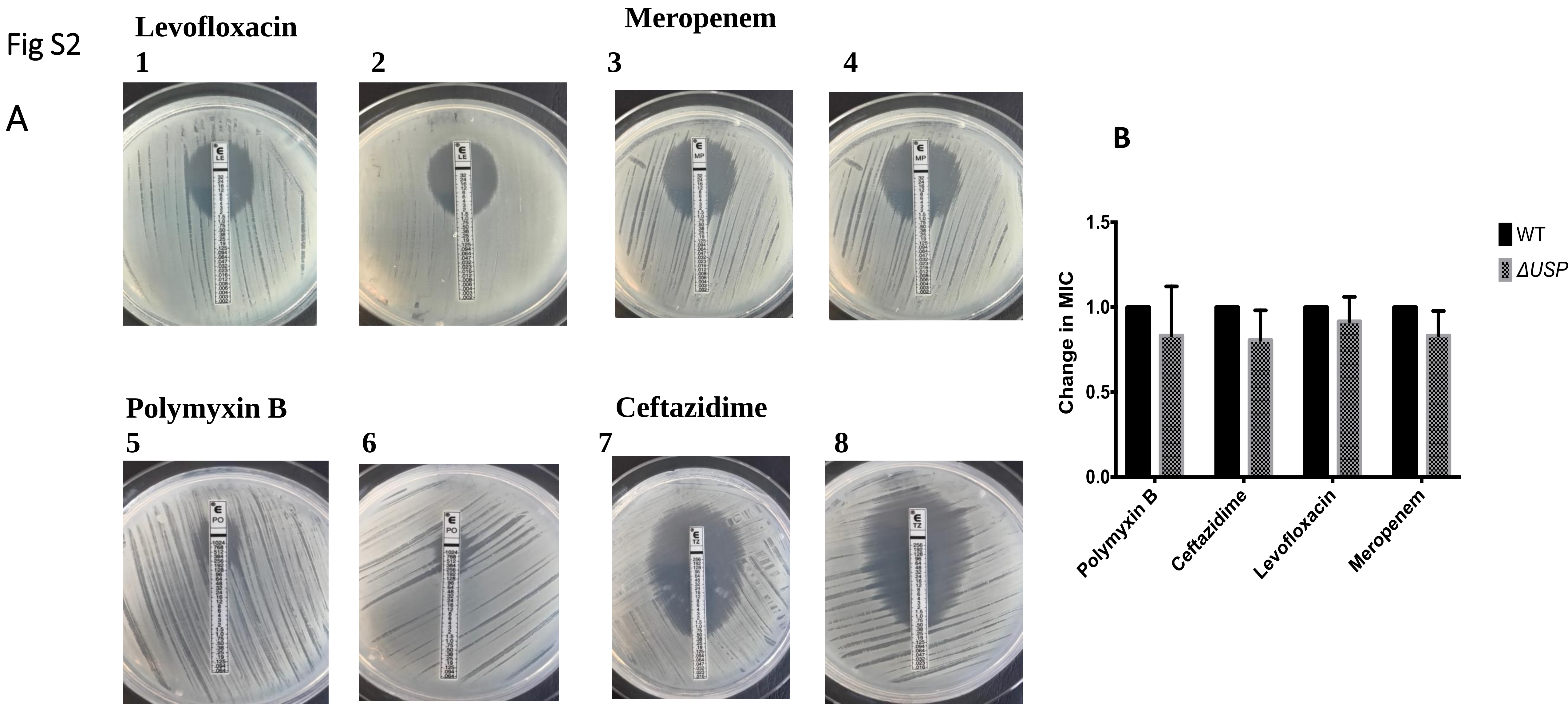

**Antibiotic susceptibility of WT (1,3,5,7)and the Δ*usp76* mutant (2,4,6,8)** using Biomerieux antibiotic ETEST® strips. Levofloxacin (1 & 2) (0.002 μg – 32 μg), Meropenem (3 & 4) (0.002 μg – 32 μg), Polymyxin B (5 & 6) (0.064 μg – 1024 μg) and Ceftazidime (7 & 8)(0.016 μg – 256 μg) performed on Mueller Hinton agar plates. **(b)** Calculated fold changes in MIC for WT and Δ*usp76* for the antibiotics using the Biomerieux ETEST® . Data shown represents the mean fold change in MIC from three independent occasions. Error bars represent the standard error

### Fig S3

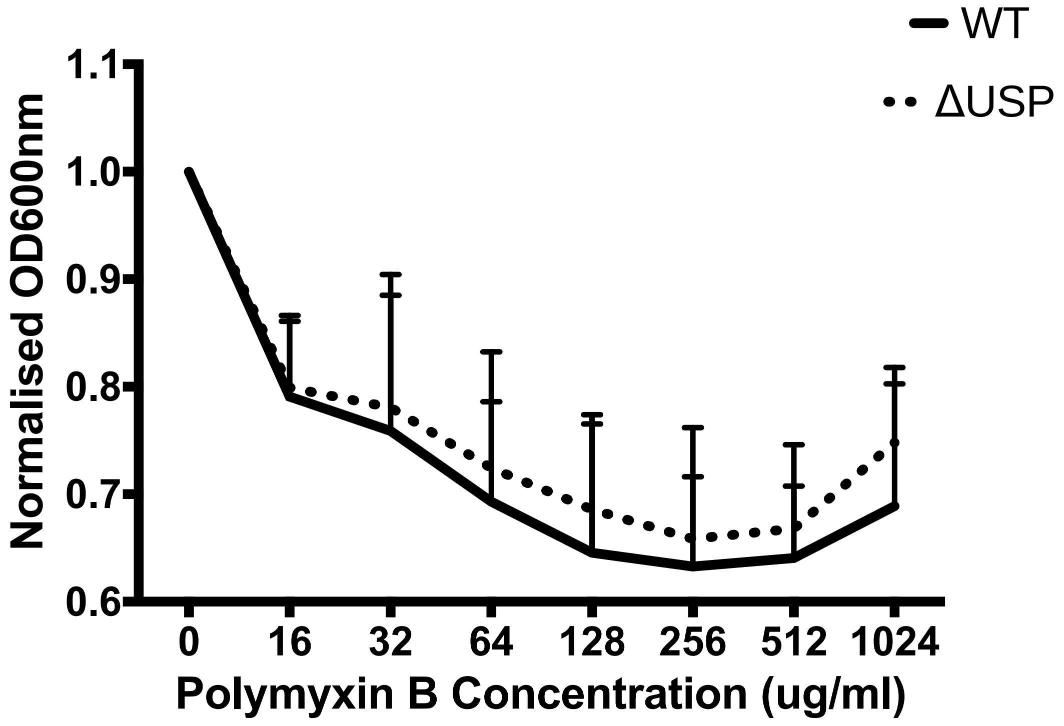

**Normalised endpoint $OD_{600nm}$ of WT and Δ*usp76* following 20 hour incubation in a range of concentrations of Polymyxin B.** Normalised endpoint $OD_{600nm}$ of WT and Δ*usp76* incubated for 20 hours in concentrations of Polymyxin B (0 – 1024 μ/ml). Data shown represents the mean $OD_{600nm}$ from three independent occasions. Error bars represent the standard error.

### Fig S4

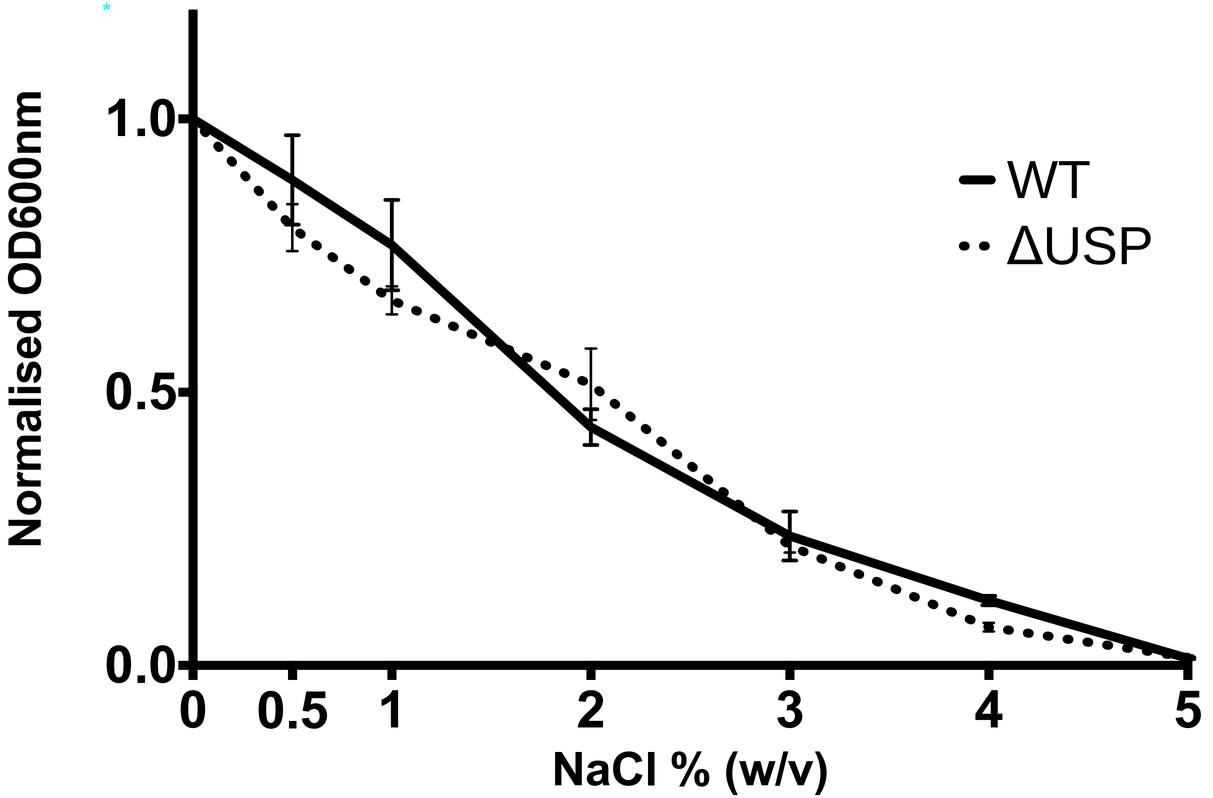

**Endpoint normalised $OD_{600nm}$ of WT K56-2 and Δ*usp76* following a 24 hours incubated with concentrations ranging from 0 – 5% W/V NaCl.** Normalised endpoint absorbance values measured at 24 hours incubation of WT K56-2 and Δ*usp76* at $37^0$C in various concentrations of NaCl ranging from 0 – 5% W/V. Values displayed have been normalised to 0% W/V NaCl (LB only with culture) and represent the average of 3 independent occasions. Statistical analysis determined by ANOVA. ($p = 0.72$).

### Fig S5

a b

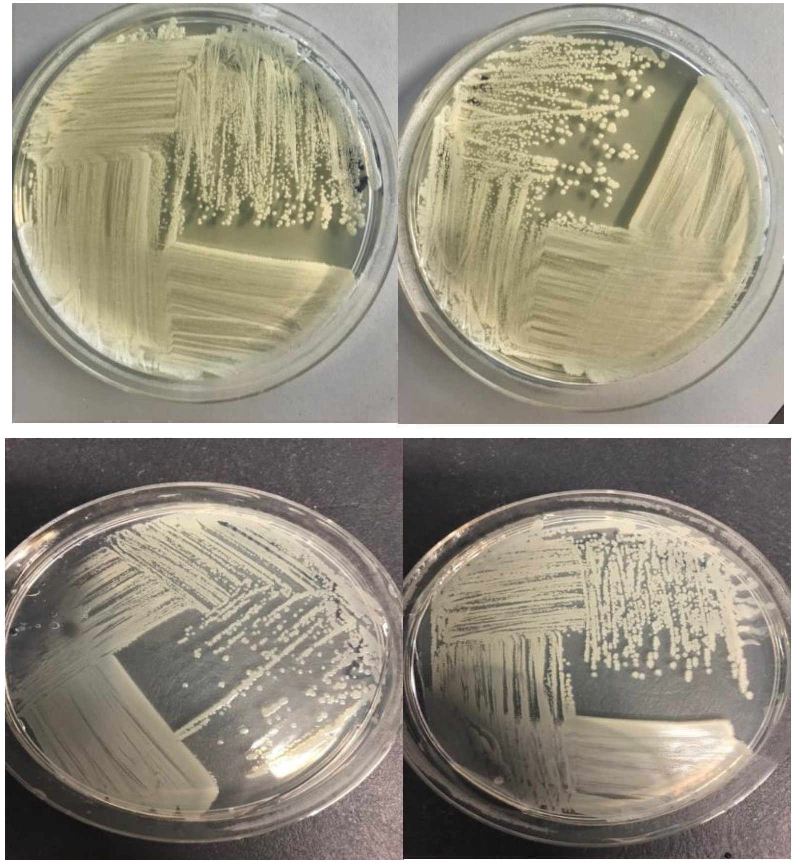

c d

**Representative images of mucoid phenotype of WT and $\Delta$*usp76* in normal oxygen conditions and at 6% $O_2$.** EPS production of WT (a) and $\Delta$*usp76* (b) incubated on YEM agar for 5 days at 37$^{o}$C in either normal oxygen conditions (A &B) or 6% $O_2$ (c & d). Images represent mucoid phenotype observed on one of three independent occasions.

**Fig S6**

**A**

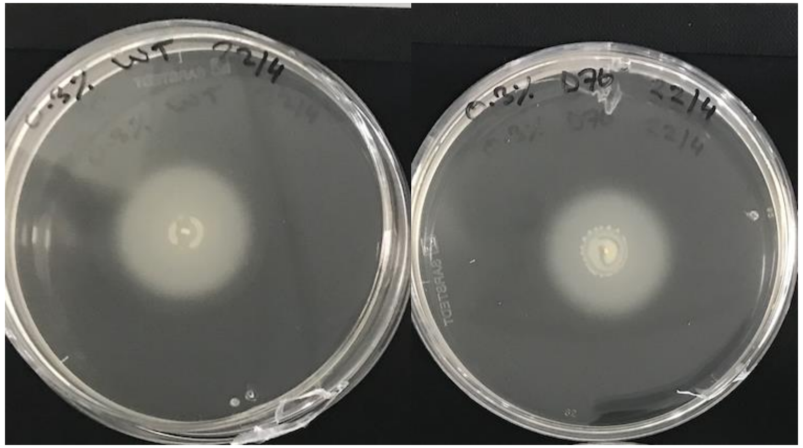

**B**

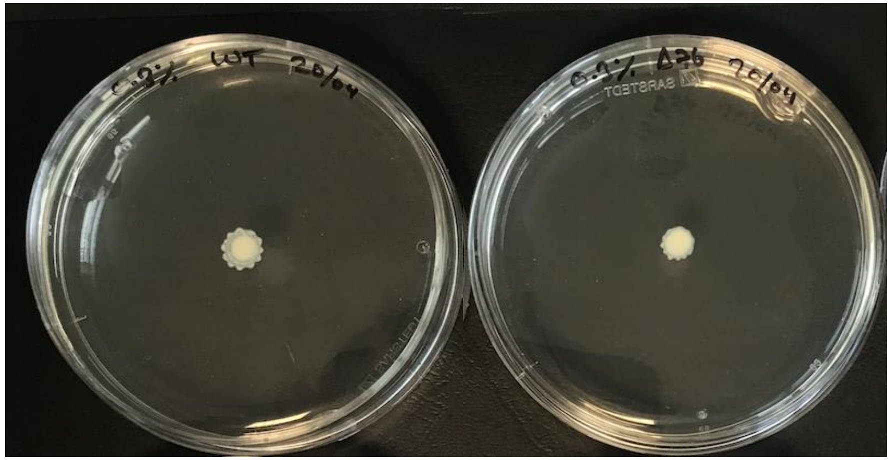

**C**

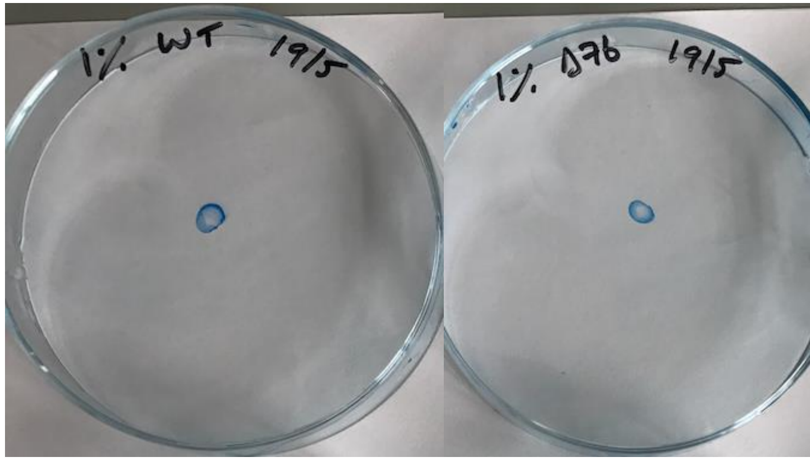

**Motility of WT and Δ*usp76*.**

**A) Swimming of strains on 0.3% w/v LB agar.** Representative images of 3 independent occasions of the assessment of swimming of WT and Δ*usp76* on 0.3% w/v LB agar incubated for 16 – 18 hours.

**B) Representative images of the assessment of swarming of WT and Δ*usp76* on 0.5% w/v LB agar.** Representative images of 3 independent occasions of the assessment of swarming of WT and Δ*usp76* on 0.5% w/v LB agar incubated for 16 – 18 hours.

**C) Representative images of the assessment of twitching of WT and Δ*usp76* on 1.0% w/v LB agar.** Representative images of 3 independent occasions of the assessment of twitching of WT and Δ*usp76* stained with Coomassie blue following incubation on 1.0% w/v LB agar for 16 – 18 hours.

**Fig S7**

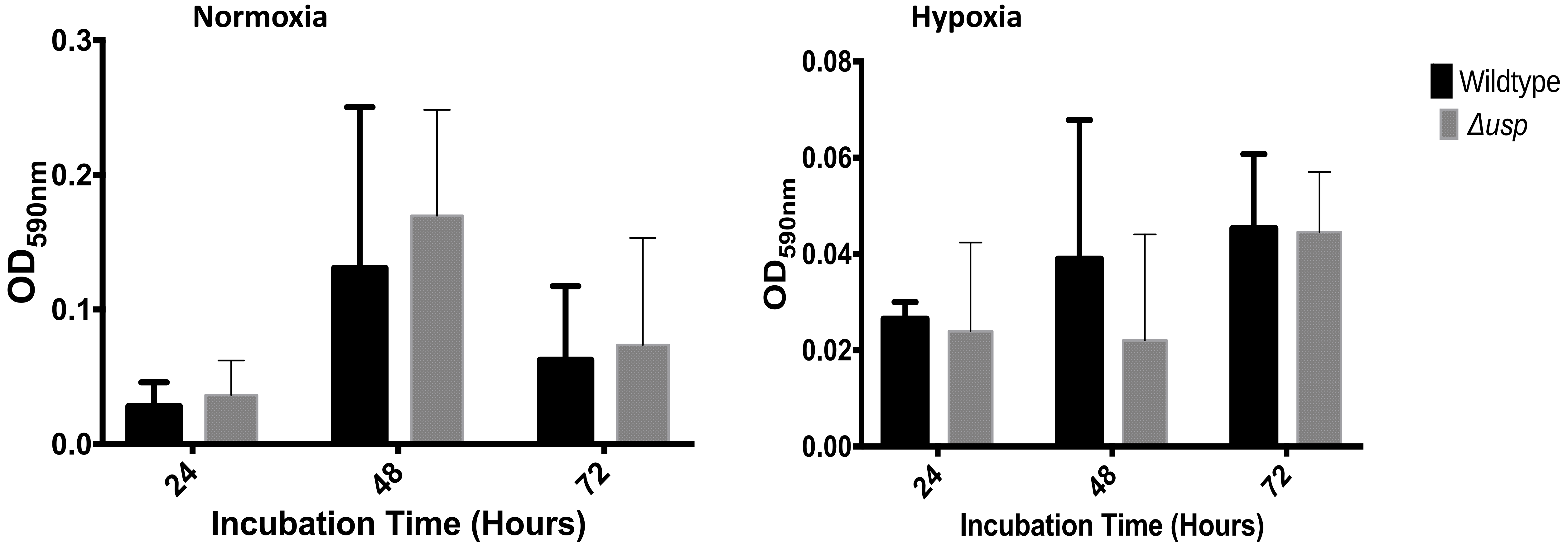

**Determination of biofilm formation of WT and Δ*usp76* mutant over a period of 72 hours in normoxic and hypoxic conditions.** Average biofilm formation for the WT and the Δ*usp76* mutant strain as determined by measuring $OD_{590nm}$ of resolubilised, crystal violet stained biofilms in normal oxygen (A) and hypoxic (6% $O_2$) (B) conditions measured every 24 hours for 72 hours. Data represents the mean $OD_{590nm}$ from three independent occasions. Error bars represent the standard error of each mean.

#### Fig S8

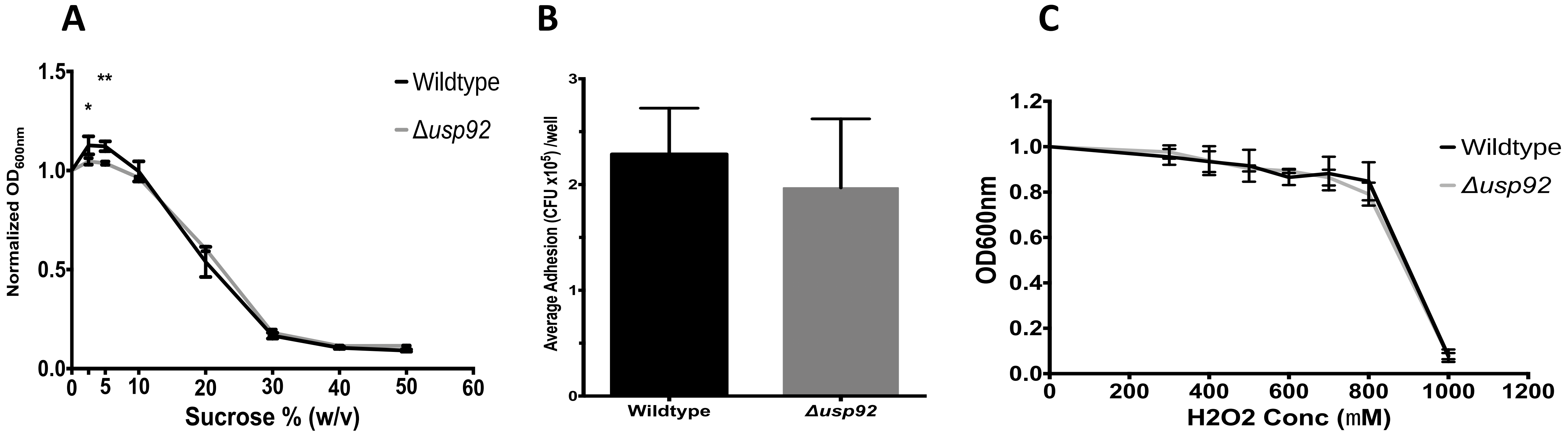

A. Average endpoint absorbance values of WT and the $\Delta$*usp92* mutant following incubation with concentrations of sucrose (0 – 50% w/v) for 24 hours at $37^{\circ}$C normalised to LB only.
B. Attachment of WT and $\Delta$*usp92* mutant CF lung epithelial cells CFBE41o- as determined by microbial CFU attached per well.
C. Average endpoint absorbance values of WT and $\Delta$*usp92* mutant following incubation with concentrations of $H_2O_2$ (0 - 1000 $\mu$M) for 20 hours. Data shown represents the mean respective values from three independent occasions. Error bars represent the standard error of the mean.

*Statistically significant difference relative to the WT as determined by two way ANOVA ($p<0.01$).

#### Fig S9

A

i

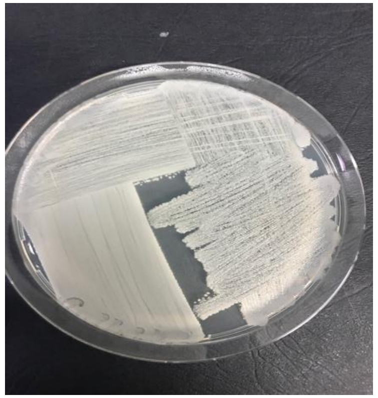

ii

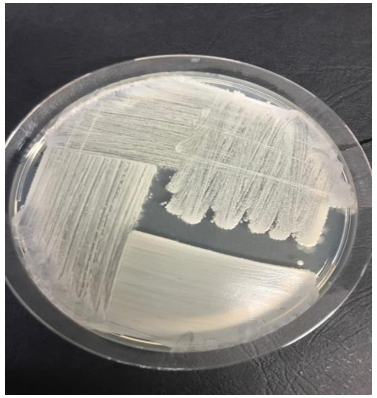

B

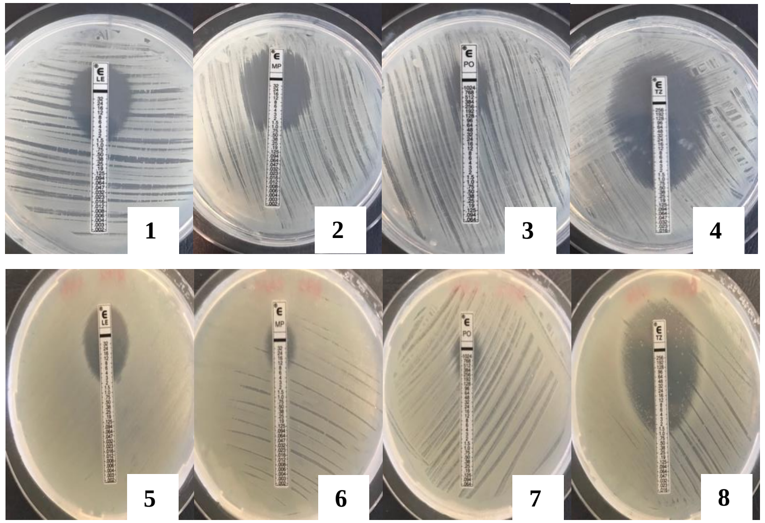

C

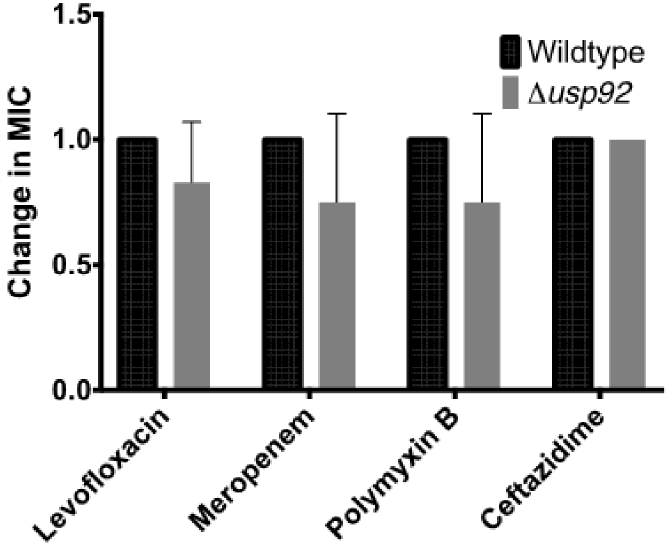

**Characterisation of the *Δusp92* mutant strain to determine the role of the BCAM0292 gene in mucoidy A) and antibiotic susceptibility (B, C).**

A. Representative images of the EPS production of WT (**i**) and the Δ*usp92* mutant (**ii**) incubated on YEM agar for five days at 37$^{o}$C in normal oxygen conditions (Images are representative of three independent occasions.

B. Antibiotic susceptibility of WT (1 to 4) and the Δ*usp92* mutant (5 to 8) using Biomerieux antibiotic ETEST® strips. Levofloxacin (1 & 5) (0.002 μg – 32 μg), Meropenem (2 & 6) (0.002 μg – 32 μg), Polymyxin B (3 & 7) (0.064 μg – 1024 μg) and Ceftazidime (4 & 8)(0.016 μg – 256 μg) performed on Mueller Hinton agar plates.

C. The mean fold change of MIC in the WT Δ*usp92* mutant in three independent experiments.
